# Antigen-specific age-related memory CD8 T cells induce and track Alzheimer’s-like neurodegeneration

**DOI:** 10.1101/2024.01.22.576704

**Authors:** Akanksha Panwar, Altan Rentsendorj, Michelle Jhun, Robert M. Cohen, Ryan Cordner, Nicole Gull, Robert N. Pechnick, Gretchen Duvall, Armen Mardiros, David Golchian, Hannah Schubloom, Lee-Way Jin, Debby Van Dam, Yannick Vermeiren, Hans De Reu, Peter Paul De Deyn, Jevgenij A. Raskatov, Keith L. Black, Dwain K. Irvin, Brian A. Williams, Christopher J. Wheeler

## Abstract

Cerebral (Aβ) plaque and (pTau) tangle deposition are hallmarks of Alzheimer’s disease (AD), yet are insufficient to confer complete AD-like neurodegeneration experimentally. Factors acting upstream of Aβ/pTau in AD remain unknown, but their identification could enable earlier diagnosis and more effective treatments. T cell abnormalities are emerging AD hallmarks, and CD8 T cells were recently found to mediate neurodegeneration downstream of tangle deposition in hereditary neurodegeneration models. The precise impact of T cells downstream of Aβ/fibrillar pTau, however, appears to vary depending on the animal model used. Our prior work suggested that antigen-specific memory CD8 T (“^hi^T”) cells act upstream of Aβ/pTau after brain injury. Here we examine whether ^hi^T cells influence sporadic AD-like pathophysiology upstream of Aβ/pTau. Examining neuropathology, gene expression, and behavior in our ^hi^T mouse model we show that CD8 T cells induce plaque and tangle-like deposition, modulate AD-related genes, and ultimately result in progressive neurodegeneration with both gross and fine features of sporadic human AD. T cells required Perforin to initiate this pathophysiology, and IFNγ for most gene expression changes and progression to more widespread neurodegenerative disease. Analogous antigen-specific memory CD8 T cells were significantly elevated in the brains of human AD patients, and their loss from blood corresponded to sporadic AD and related cognitive decline better than plasma pTau-217, a promising AD biomarker candidate. Our work is the first to identify an age-related factor acting upstream of Aβ/pTau to initiate AD-like pathophysiology, the mechanisms promoting its pathogenicity, and its relevance to human sporadic AD.

**Significance Statement:** This study changes our view of Alzheimer’s Disease (AD) initiation and progression. Mutations promoting cerebral beta-amyloid (Aβ) deposition guarantee rare genetic forms of AD. Thus, the prevailing hypothesis has been that Aβ is central to initiation and progression of all AD, despite contrary animal and patient evidence. We show that age-related T cells generate neurodegeneration with compelling features of AD in mice, with distinct T cell functions required for pathological initiation and neurodegenerative progression. Knowledge from these mice was applied to successfully predict previously unknown features of human AD and generate novel tools for its clinical management.

## Introduction

Deposits of aggregated amyloid plaques containing beta-amyloid (Aβ) and neurofibrillary tangles (NFTs) comprised of hyper-phosphorylated tau (pTau) are the primary neuropathological hallmarks of Alzheimer’s disease (AD). Aβ deposition precedes that of NFTs in all forms of AD, and rare inherited forms of the disease are due to mutated (AD-mut) genes that promote the deposition of toxic Aβ in brain. Thus, a prominent hypothesis is that Aβ is central to initiation and progression of all AD. While Aβ deposition clearly plays an important role, evidence that it is sufficient to initiate and/or maintain AD pathogenesis is lacking. For example, laboratory animals expressing one or even several human AD-mut transgenes that guarantee AD in their human carriers fail to exhibit either robust neurodegeneration or NFTs (1-3). Moreover, targeting Aβ deposition has resulted in hundreds of failed clinical trials, with modest and sometimes controversial clinical efficacy demonstrated in a few trials only recently (4). Finally, up to a third of elderly individuals possess Aβ deposition sufficient for AD diagnosis, yet are cognitively normal and may remain so for life (5). These findings suggest that Aβ requires distinct co-factors to initiate AD. The search for such factors, with occasional exceptions (6), has primarily focused on processes downstream of Aβ. A recent finding, however, documented that a patient harboring a deterministic AD-mut gene normally guaranteeing neurodegeneration by age 45, exhibited no clinical symptoms into their 70s (7). This protection was afforded by a second germline mutation in the ApoE3 gene that disrupts lipid metabolism (8), which highlights the possibility that the pathogenic effects of Aβ may be dominantly inhibited by pre-existing physiological processes upstream of Aβ. While the precise nature of such processes remains unknown, identifying them could allow earlier diagnosis and lead to more effective AD treatments.

Among Aβ-independent factors potentially contributing to AD pathogenesis, CD8 T cell abnormalities have recently emerged as some of the most intriguing. Recent studies have verified and extended prior reports of T cell abnormalities in AD-related conditions, including resident memory CD8 T (T_RM_) and other memory T cell accumulation in CSF and/or brain of aging and AD-afflicted individuals (9, 10). CD8 T cells within CSF also dysregulate chemokine signaling when cognitive impairment is evident, regulate neuronal and synaptic gene expression when they accumulate in hippocampus, and promote neuroinflammation and cognitive decline, in transgenic mice (11-13). Moreover, T cells increase in areas of AD-Tg mouse brain and human AD brain with tau neuropathology (11), which correlates with cognitive symptoms in AD (14, 15). Most surprisingly, CD8 T cells together with microglia and interferon-gamma (IFNγ), were recently shown to mediate neurodegeneration in tau-transgenic mice (16). Finally, the most recent study on CD8 T cells in AD-Tg mice presented evidence that they actually dampen Aβ/pTau pathology through CXCL16-CXCR6 intercellular communication between T cells and microglia (17). Thus, the role of CD8 T cells appears variable, at least in in transgenic rodents. Moreover, T cell changes in AD are often presumed to follow Aβ deposition. For example, in all of the studies above T cell abnormalities were examined after AD pathology was established. The potential role of CD8 T cells upstream of Aβ (or tau) has thus not been examined. Nevertheless, the potential relationship of tissue-resident CD8 T cells in particular to ApoE-mediated AD risk and/or protection makes them especially intriguing in this regard (18).

In order to examine the effect of aging T cells upstream of Aβ/pTau pathology, we recently developed the “^hi^T” model of rapid CD8 T cell aging in young mice. In this model, age-related homeostatic CD8 T cell expansion is mimicked on a much faster time scale (19). This results in morphological, functional, and genotypic changes in homeostatically induced (“hi”) T cells similar to those occurring with age. These changes include human-like predominance of CD103^+^ memory CD8 T cells in the general circulation, and accumulation of an antigen(Amyloid Precursor Protein; APP)-specific age-related subpopulation of T cells in brain. Transfer of these age related CD8 T cells from ^hi^T mice into wild-type B6 mice also increased human-like amyloid and fibrillar pTau in brain after percussive cerebral injury, as well as neuronal marker loss regardless of injury (19). This suggested that age-related CD8 T cells act upstream to promote both Aβ and fibrillar pTau increases in a non-AD setting. Here, we examine the possibility that age-related CD8 T cells are directly related to events upstream of Aβ, and possibly other aspects of AD pathophysiology, by first interrogating proteinopathy, neurodegeneration, and behavior in the ^hi^T mouse model. We then examined mechanisms whereby ^hi^T cells mediate these pathological characteristics using T cells from functional knockout donors as well as RNAseq analysis. Finally, we examined the clinical relevance of antigen-specific and parental CD8 T cell populations to human AD using flow cytometric, tissue staining, and protein analysis in human blood and brain.

## Results

### Creation of ^hi^T mouse

PBS, or CD8 T cells from wild-type donors were injected into young (6-8 week) B6.Foxn1 hosts (PBS and wt-CD8 groups, respectively). Limited analyses of young B6.Foxn1 hosts injected with wild-type CD4 T cells or CD4 plus CD8 T cells together (wt-CD4 and wt-CD8+CD4 groups, respectively), were also performed (Supplementary Materials fig. S1A). Females were exclusively used to avoid male-specific autoimmune disease dynamics, some of which are associated with CD8 T cell homeostatic expansion, in related mouse strains (20, 21). Injection results in rapid homeostatic expansion of the donor cells in the circulation, similar to the gradual homeostatic expansion of CD8 T cells that occurs with aging in humans. The donor T cells then acquire age-related resident memory-like phenotype, as well as age-related genotype and function as they expand. CD8 T cells reactive to a non-Aβ epitope on Amyloid Precursor Protein (APP) then selectively accumulate in brain of these “^hi^T” (for “homeostatically induced T cell”) mice (19). Circulating APP-reactive CD8 T_RM_ cells were substantially expanded in ^hi^T mice (Supplementary Materials fig. S1B), similar to the reported expansion of age-related CD8 T_RM_ cells in aging humans (22).

### Aβ and neurofibrillary deposition

We examined Triton-soluble brain extracts as we previously described (23), to assess potentially toxic Aβ in ^hi^T mice by Western blot with ab14220 (rodent-specific Aβ). This revealed relative upregulation of high molecular weight species presumed to be Amyloid Precursor Protein (APP) and its cleavage products in brains of wt-CD8 group ^hi^T mice 3-10 weeks after injection (Fig. 1). Additional isoform- and species-specific antibodies in ELISA analysis revealed that endogenous Aβ1-40 but not Aβ1-42 was relatively increased in wt-CD8 group mice 10 weeks post-injection, with Western blots and tissue staining using 4G8 antibody (rodent and human Aβ/APP) confirming prominent amyloidosis in cortex and hippocampus (Fig. 1A-C; Supplementary Materials fig. S2A). By 6 months post-injection, wt-CD8 and wt-CD8+CD4 groups exhibited increased Aβ deposition in brain vasculature, whereas wt-CD4 group did not (Fig. 1D; Supplementary Materials fig. S2A-F) (24), further suggesting CD8 T cells are necessary and sufficient to promote amyloidosis. Aβ plaques in wt-CD8 group brains 15 months post-injection exhibited typical morphology, but exhibited small compact cores at best, were chiefly detergent-soluble (Supplementary Materials fig. S3), and were minimally stained by curcumin or ThioS (Fig. 1C).

**Fig. 1:**
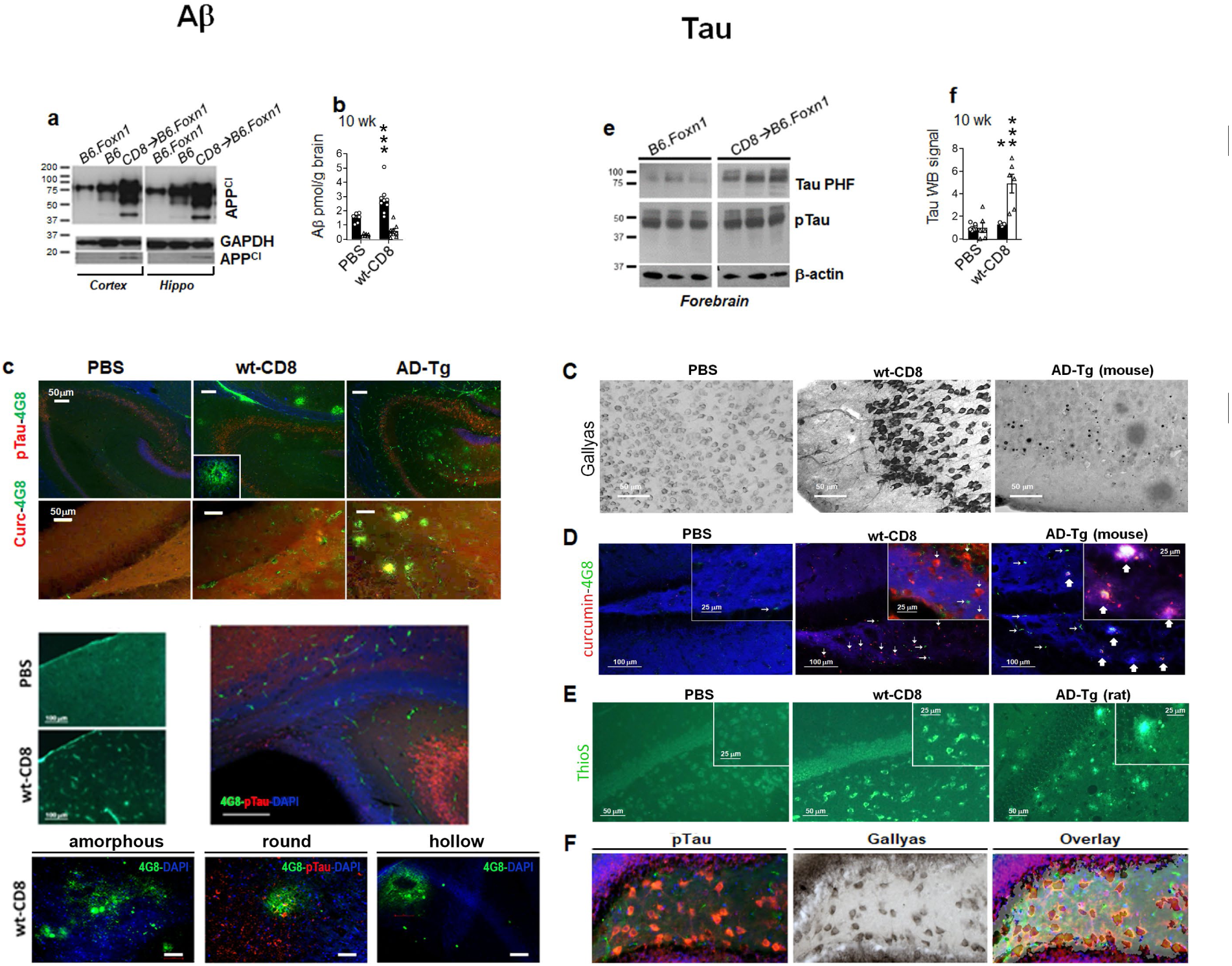
Amyloid pathology. Western blots of APP cleavage products (APP^Cl^) in cortex, hippocampus 3 wk after injection (◊) of recipients **(A)**. **B-G** depict B6.Foxn1 recipients, 15 mos post-injection unless otherwise indicated. Forebrain Aβ1-40/42 ELISA **(B)**. Plaques ± pTau/curcumin staining **(C)**, and compiled 4G8 burden in entorhinal (Ent)/cingulate (Cng) cortex, and hippocampus (Hippo) **(D)**. Measurement bars on immunofluorescent images in **C** are all 50μm, and in B are all 20 μm in length.

Detergent-soluble phospho-tau (pTau) was slightly (30%) but significantly increased at 10 weeks post-injection in wt-CD8 group forebrain, while pTau paired helical filaments (PHFs, which mature to form NFTs in AD) were increased nearly 5-fold on Western blots (Fig. 2A). Starting at 6 months post-injection, fibril-staining reagents (Gallyas silver, curcumin, and Thio-S), each stained cellular inclusions within wt-CD8 group hippocampus, with no similar structures seen in simultaneously stained AD-transgenic Tg2576 (AD-Tg) mice and/or rats (Fig. 2C-E) (23). Tissue immunofluorescence for pTau prior to Gallyas staining revealed these structures were derived from pTau^+^ neurons with intact nuclei, and that silver staining was superimposable with that of pTau but distinct from Aβ (Fig. 2F, G). These data suggest that the predominant T cells in hiT mice (which resemble resident-memory phenotype CD8 T cells or CD8 T_RM_) promote coordinated and robust deposition of Aβ and fibrillar NFT-like inclusions in mouse brain.

**Fig. 2:**
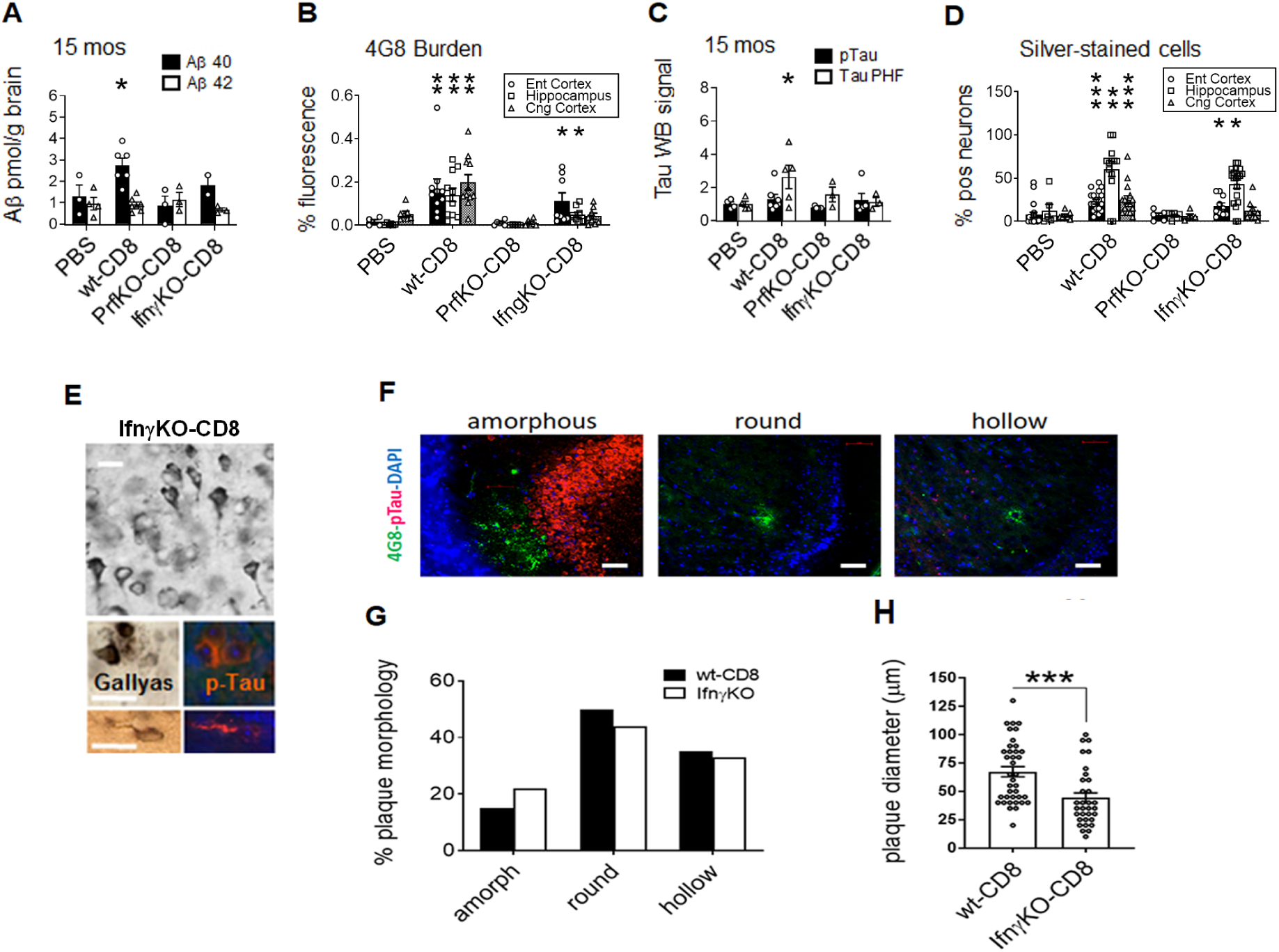
Tau pathology. Western blots **(A)**, and compiled pTau and PHF signal (pS199/202 antibody [Invitrogen] was used for pTau in Western blots and tissue staining; Phospho-PHF-tau pSer202+Thr205 Antibody [AT8] was used for Tau PHF in Western blots)**(B)**. Gallyas/silver-stained cells in ^hi^T_RM_ groups and 18-month-old Tg2576 (AD-Tg) mice, with sequential pTau→Gallyas stains inset **(C)**. Hippocampal sections from the indicated groups (all B6.Foxn1 recipients, except AD-Tg = Tg2576 mice), were stained with 4G8 (Aβ) and curcumin, 6 months after i.v. control/cell injection, or at 14 months of age for AD-Tg **(D)**. Right-facing arrows highlight Aβ deposits with no curcumin co-staining. Up-facing arrows depict co-localized Aβ and curcumin deposits, representing mature neuritic plaques. Down-facing arrows highlight curcumin^+^ structures with no Aβ co-staining, i.e., non-amyloid fibrillar deposits. No DAPI was used; blue background is provided for anatomical context. ThioS staining of PBS and wt-CD8 group B6.Foxn1 ^hi^T recipients 6 months after control/cell injection, and 20 month-old AD-Transgenic (Tg) rat dentate gyrus **(E)**. Rat AD-Tg brain failed to stain with ThioS in our hands, despite prior reports of NFT content (Cohen et al., 2013). Measurement bars on immunofluorescent images are 25μm in length unless otherwise specified. Sequential Gallyas → pTau staining of neurons in wt-CD8 hippocampus **(F)** and IfnγKO-CD8 cortex **(G)** and Gallyas^+^ neurons from IfnγKO-CD8 group cortex **(G)**. Compiled Gallyas^+^ neurons in all B6.Foxn1 ^hi^T recipients 15 months after control/cell injection **(H)**. Plots depict averages ± SEM. \**P* < 0.05, \*\**P* < 0.01, \*\*\**P* < 0.005 by 2-sided T-test, relative to PBS group.

### Immune & neuroinflammatory infiltration

Additional images support our previous findings that CD8 T cell infiltration (Supplementary Materials fig. S5A-C), astrogliosis (Supplementary Materials fig. S5D,E), and microgliosis (Supplementary Materials fig. S5F,G), are exclusively seen in wt-CD8 group mice (19). Cerebral plaques in these animals were closely associated with microglia and astrocytes, as is common in human AD (Supplementary Materials fig. S5H,I). Aβ plaque burden correlated most strongly with hippocampal CD8 T cell levels rather than astrocytic or microglial levels (Supplementary Materials fig. S5J-L), suggesting a unique relationship between CD8 T cells and amyloid pathology.

### Neuronal loss & cerebral atrophy

To determine if neurodegeneration was evident in ^hi^T mice, we stained and counted neurons positive for the neuron specific nuclear protein, NeuN, in CA1, CA2, and CA3 of hippocampus, assessed brain mass, and quantified NeuN and synaptic protein signals on Western blots. Loss of NeuN^+^ cells in wt-CD8 group mice was visually apparent in hippocampal immunostains, and was verified by NeuN^+^ cell counts at 15 months post-injection (Fig. 3A-C). Loss of brain mass in wt-CD8 group progressed from 5% at 6 months, to 10% 15 months post-injection (Fig. 3D), comparable to terminal brain atrophy in human AD (25). Western blots of brains 15 months post-injection confirmed proportional decreases in NeuN, the post-synaptic submembrane protein, Drebrin, and the pre-synaptic vesicle protein, Synaptophysin (Fig. 3E, F). Loss in brain mass correlated with decreased NeuN within PBS and CD8 injection groups, establishing a direct relationship between brain atrophy and neuronal loss (Fig. 3G). No significant brain loss was observed in wt-CD4 group, but was evident in wt-CD8+CD4 group, suggesting that CD4 T cells fail to either mediate neurodegeneration or modulate its induction by CD8 T cells (Supplementary Materials fig. S6A).

**Fig. 3:**
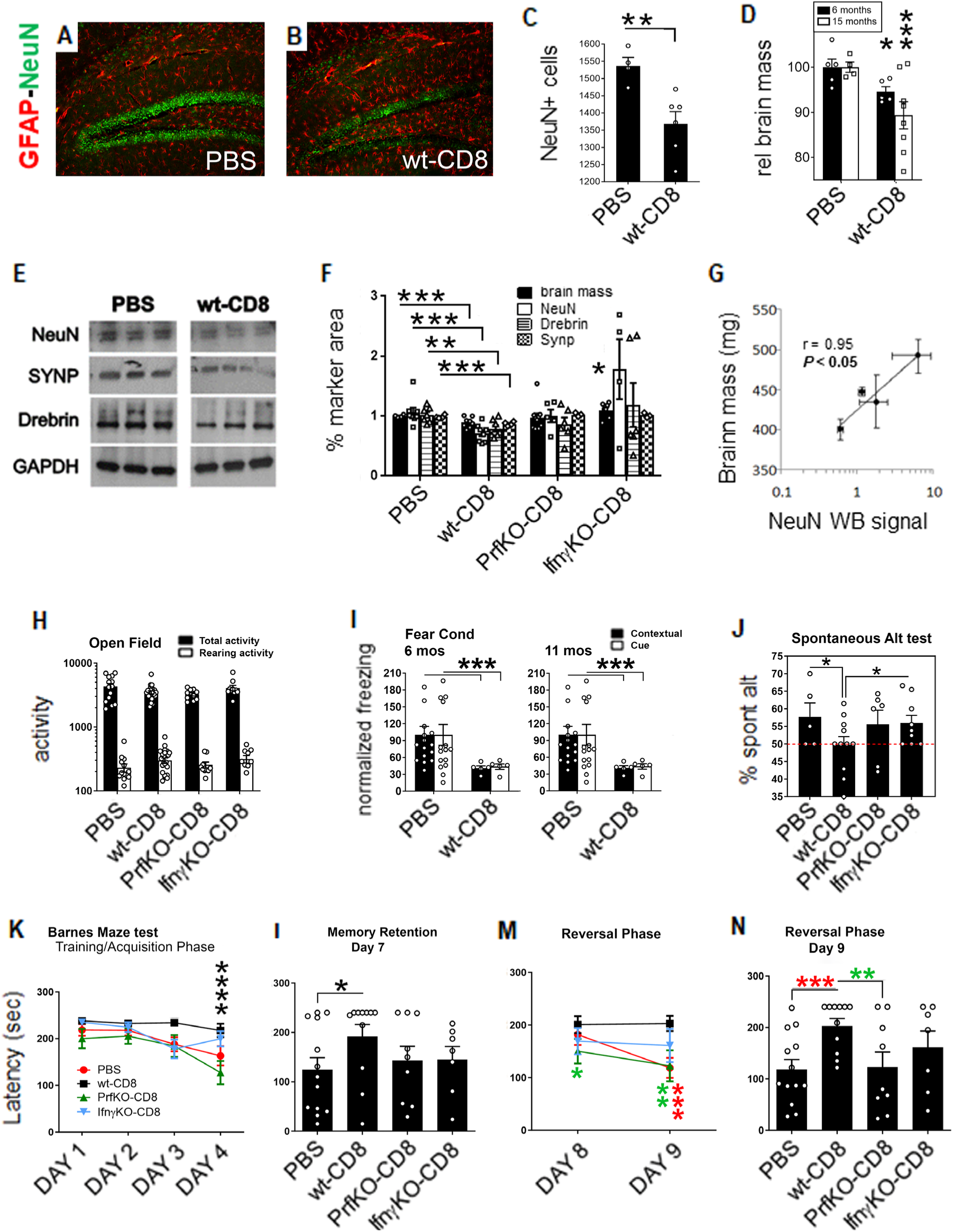
Neurodegeneration and cognition in nude mice harboring ^hi^T cells. Cell/control recipients in all panels are B6.Foxn1 exclusively. NeuN and GFAP staining **(a, b)**, and cell counts in hippocampus, 15 mos after cell/control injection **(c)**. Brain atrophy over time in PBS and wt-CD8 groups (mass normalized to PBS controls at each time point; **d)**. Representative forebrain Westerns (**e**), and GAPDH-normalized NeuN, Drebrin, and Synaptophysin Western signals (**f**). Correlation of NeuN with brain weight (**g)**. Plots depict averages ± SEM. \**P* < 0.05, \*\**P* < 0.01, \*\*\**P* < 0.005 by 2-sided T-test, relative to PBS group. Representative Open Field test at 13 mos **(h)**. Fear Conditioning over time **(i)**, and Spontaneous Alternation Behavior at 12 months **(j)**. Barnes Maze learning/training **(k)**, retention **(l)**, and reversal **(m, n)** phases, at 14 mos (black, colored symbols = *P* relative to PBS, wt-CD8, respectively). Plots depict averages ± SEM. \**P* < 0.05, \*\**P* < 0.01, \*\*\**P* < 0.005 by 2-sided ANOVA (panel **k**) or 2-sided T-test (all others), relative to PBS group unless otherwise indicated.

### Severe cognitive impairment

We assessed cognitive performance in ^hi^T mice using 3 independent tests for learning and memory: contextual and cued Fear Conditioning (FC), Spontaneous Alternation Behavior (SAB) in the Y-maze, and Barnes Maze (BM) performance. Open Field testing (OFT) was performed prior to each behavioral test (Fig, 3H; Supplementary Materials fig. S7A) to rule out motor dysfunction such as that characterizing multiple sclerosis.

Significant motor deficits occurred with age, but did not distinguish control from treatment groups (Supplementary Materials fig. S7A). In contrast, contextual FC was reduced in wt-CD8 group relative to PBS controls 6 months after T cell injection, with both contextual and cued learning impaired at 11 months (Fig 3I). These results suggest that CD8 T cells in ^hi^T mice alone mediate early damage to hippocampus (required for contextual FC), with additional later damage to amygdala (involved in cued FC), mirroring the typical pattern of cognitive decline in human AD (26). Intriguingly, only cued learning was impaired in wt-CD8+CD4 but not wt-CD4 group mice at the earlier time point (Supplementary Materials fig. S6B), suggesting that CD4 T cells modulate whether hippocampus or amygdala is damaged first by ^hi^T_RM_, but not their overall neurodegenerative impact.

SAB is a well established assay for hippocampal integrity (27-29), based on the preference of mice to alternately explore two alleys. The lowest possible score (50%) indicates random alley choice, due to either no working memory of the previous alley entered, or complete lack of preference. SAB testing 12 months post-injection revealed a score of 55-56% in PBS controls, comparable to wild-type mice (30), but was reduced to 50% in wt-CD8 group (Fig. 3J), consistent with no working memory.

Barnes Maze testing was performed at 14 months post-injection, and represents a more focused measure of hippocampus-dependent learning and memory, that is further not confounded by decreased ability to swim, or stress-induced behaviors caused by swim tests (31, 32). In contrast to all other groups, wt-CD8 mice showed no ability to learn the maze over the initial 4-day training period (Fig. 3K). Given this initial deficit, wt-CD8 mice were also profoundly impaired on subsequent memory retention and reversal phases of the maze (Fig. 3L-N).

Contextual FC at 6 and 11 months correlated with brain mass (Supplementary Materials fig. S7B), as did latency to solve the Barnes Maze (Supplementary Materials fig. S7C), further underscoring the relationship of cognitive decline to physical neurodegeneration. Poor performance of ^hi^T mice on Barnes Maze (total latency below median = BM^lo^) also exhibited significant association with increased pTau PHFs exclusively (Supplementary Materials fig. S7D), but not with any form of Aβ1-40/42 or detergent-soluble pTau (Supplementary Materials fig. S7E,F).

Taken together, these three independent tests suggest that fully functional ^hi^T_RM_ mediate severe, progressive impairment of hippocampus-dependent learning and memory independent of locomotor activity, and that the ^hi^T model mirrors patterns of cognitive loss and association with cerebral pathology seen in clinical AD (33).

### Mechanisms of ^hi^T cell effects

To examine mechanisms of ^hi^T cell mediated neuropathology, we injected PBS, or CD8 T cells from wild-type, Perforin 1-deficient, or IFNγ-deficient donors into young (6-8 week) B6.Foxn1 hosts (PBS, wt-CD8, PrfKO-CD8, and IfnγKO-CD8 groups, respectively), and analyzed neuropathology by ELISA or Western blot, and tissue staining. In marked contrast to wt-CD8 or IfnγKO-CD8, PrfKO-CD8 brains exhibited no increase in Aβ, plaques, pTau, PHFs, or NFT-like inclusions in any region. CD8 T cells expanded in circulation of all B6.Foxn1 recipients, with those reactive to a non-Aβ epitope on Amyloid Precursor Protein (APP) selectively accumulating in brain (19). At 15 months post-injection, Aβ1-40 was significantly elevated only in wt-CD8 group brain by ELISA (Fig. 3A). Nevertheless, Aβ plaques and silver-stained neurons were evident within entorhinal cortex and hippocampus by tissue immunofluorescence in both wt-CD8 and IfnγKO-CD8 groups, and the latter were derived from pTau^+^ neurons by sequential staining (Fig. 1B, E, F). Aβ plaques in these two groups were of similar morphology (Fig. 3F, G), but those in IfnγKO-CD8 brains were significantly smaller (Fig. 1H), and appeared only in entorhinal cortex and hippocampus but not in cingulate cortex as in the wt-CD8 group (Fig. 1B). Tau PHFs were also only elevated in wt-CD8 brains at 15 months by Western blot, but Gallyas tissue staining of IfnγKO-CD8 brains revealing NFT-like inclusions in entorhinal cortex and hippocampus, but not in cingulate cortex as in wt-CD8 group brains. These data suggest that blocking Perforin production completely prevented all signs of AD-like pathology, while blocking IFNγ production led to smaller Aβ plaques and limited distribution of NFT-like like structures reminiscent of early-stage AD **(34)**.

Neuronal and cognitive loss differences were also evident between wt-CD8 and KO-CD8 groups: neither IfnγKO-CD8 nor PrfKO-CD8 brains showed significant evidence of neuronal loss, and both were indistinguishable from PBS controls in level of Spontaneous Alternation (Fig 2J). IfnγKO-CD8 mice, however, appeared initially similar to PBS controls in the training sessions of the Barnes Maze, but became more similar to wt-CD8 group mice at many of the later stages (Fig. 2. This suggests that both IFNγ and PRF1 deficiency eliminates robust neuronal loss and either subdues cognitive decline (in IfnγKO-CD8 group) or eliminates it altogether (in PrfKO-CD8 group). The more dramatic impact of PRF1 deficiency on AD-like pathophysiology in particular is most consistent with a direct role for lytic elimination of neural cells by CD8 T cells, which requires expression of MHC I. Most neuronal lineage cells are MHC I-negative unless induced by cytokines such as IFNγ, but catecholaminergic neurons within the locus coeruleus (35) and Doublecortin(DCX)-positive neural progenitors consitutively express these molecules. To determine if such cells were eliminated early and independent of IFNγ, we examined DCX^+^ neural progenitors within the subventricular zone (SVZ) of wt-CD8 and IfnγKO-CD8 group mice.

### Reduction of Doublecortin (DCX)^+^ neurons in ^hi^T mice

CD8 T cells were observed in close proximity to DCX^+^ neural progenitors in the subventricular zone (SVZ) rostral migratory stream of wild-type mice, consistent with the possibility that CD8 T cells might directly eliminate these cells under disease conditions (Supplementary Materials fig. S8A). Accordingly, DCX^+^ neurons were also dramatically decreased in SVZ and throughout brain in wt-CD8 and IfnγKO-CD8 group ^hi^T mice (Supplementary Materials fig. S8B,C). Thus, even in the absence of more widespread neuronal loss, ^hi^T_RM_ promoted DCX^+^ promoted neural progenitor elimination. This suggests that neuronal lineage cells that express MHC I are among the earliest cells eliminated by ^hi^T_RM_, while more widespread neuronal elimination requires T cell production of the MHC I-inducing cytokine, IFNγ (36).

### RNAseq analysis on brains of ^hi^T mice

To gain further mechanistic insights into howCD8 T cells in ^hi^T mice promote AD-like pathology, we performed RNAseq analysis on mouse forebrains from experimental (wt-CD8, IfnγKO-CD8, PrfKO-CD8) and control (PBS) groups. Focusing on changes ≥10% relative to PBS, wt-CD8, IfnγKO-CD8, and PrfKO-CD8 groups exhibited significant upregulation of the T cell-specific marker, CD3ε, while wt-CD8, IfnγKO-CD8, and PrfKO-CD8 groups also altered 1947, 309, and 1901 additional transcripts, respectively (Supplementary Materials fig. S9A-I). Since this was at odds with our previous observation of no detectable CD8 T cells in PrfKO-CD8 group brain by tissue staining (19), we re-examined T cells in this group using potentially more sensitive flow cytometric analysis (Supplementary Materials fig. S9). Indeed, flow cytometry confirmed significantly elevated CD8 T cells in PrfKO-CD8 group brain relative to untreated B6.Foxn1 (Supplementary Materials fig. S9A), albeit somewhat lower relative to wild-type C57BL/6 controls (Supplementary Materials fig. S9B). Thus, all groups injected with CD8 T cells harbored elevated levels of these cells in brain, consistent with their CD8ε mRNA upregulation and providing proper context for gene expression analysis.

Two gene expression pathways were prominently altered in the wt-CD8 group exclusively: BioPlanet Alzheimer’s Disease pathway, and a group of largely overlapping pathways related to cytoplasmic ribosomal proteins and translation (Fig. 4A, B; Supplementary Materials fig. S9H). In addition, a group of largely overlapping pathways related to electron transport chain and oxidative phosphorylation was upregulated in both wt-CD8 and IfnγKO-CD8 brain (Fig. 4B; Supplementary Materials fig. S9H), and as such was uniquely associated with both early and late AD-like neuropathology. Electron transport chain- and ribosomal protein-related pathways are known to be involved in CD8 T cell-mediated effector activity as well as AD pathophysiology, and IFNγ is also known to modulate ribosomal proteins (37). Within cell type-specific genes, only neuronal genes exhibited net upregulation, whereas most non-neuronal genes were downregulated (Fig. 4C). Moreover, multiple pathways implicated in AD were affected in the wt-CD8 group (Supplementary Materials fig. S10). To further determine relevance of these modest changes to AD, we focused on 82 genes identified in prior genome-wide epidemiological studies (“GWAS genes”) (38) whose AD-associated variants often manifest as subtle expression changes in human AD. Among this GWAS subset, 14, 2, and 11 genes were significantly altered in wt-CD8, IfnγKO-CD8, and PrfKO-CD8 groups, respectively, with 8 and 1 of these also altered in wt-CD8 and IfnγKO-CD8 relative to PrfKO-CD8 (*P* = 0.034; 2-sided Fisher’s Exact Test; Fig. 4D). Known functions of GWAS genes altered in wt-CD8 and PrfKO-CD8 brains included multiple biological pathways implicated in AD (Supplementary Materials fig. S9J), whereas those altered in IfnγKO-CD8 brains covered a more restricted subset involved in Aβ/Tau pathology and inflammation/immunity. This supports a model in which Perforin controls a critical disease-initiating event reflected by electron transport chain gene modulation, whereas IFNγ modulates most other gene expression risk factors that may worsen disease pace and/or severity (Fig. 4E).

**Fig. 4:**
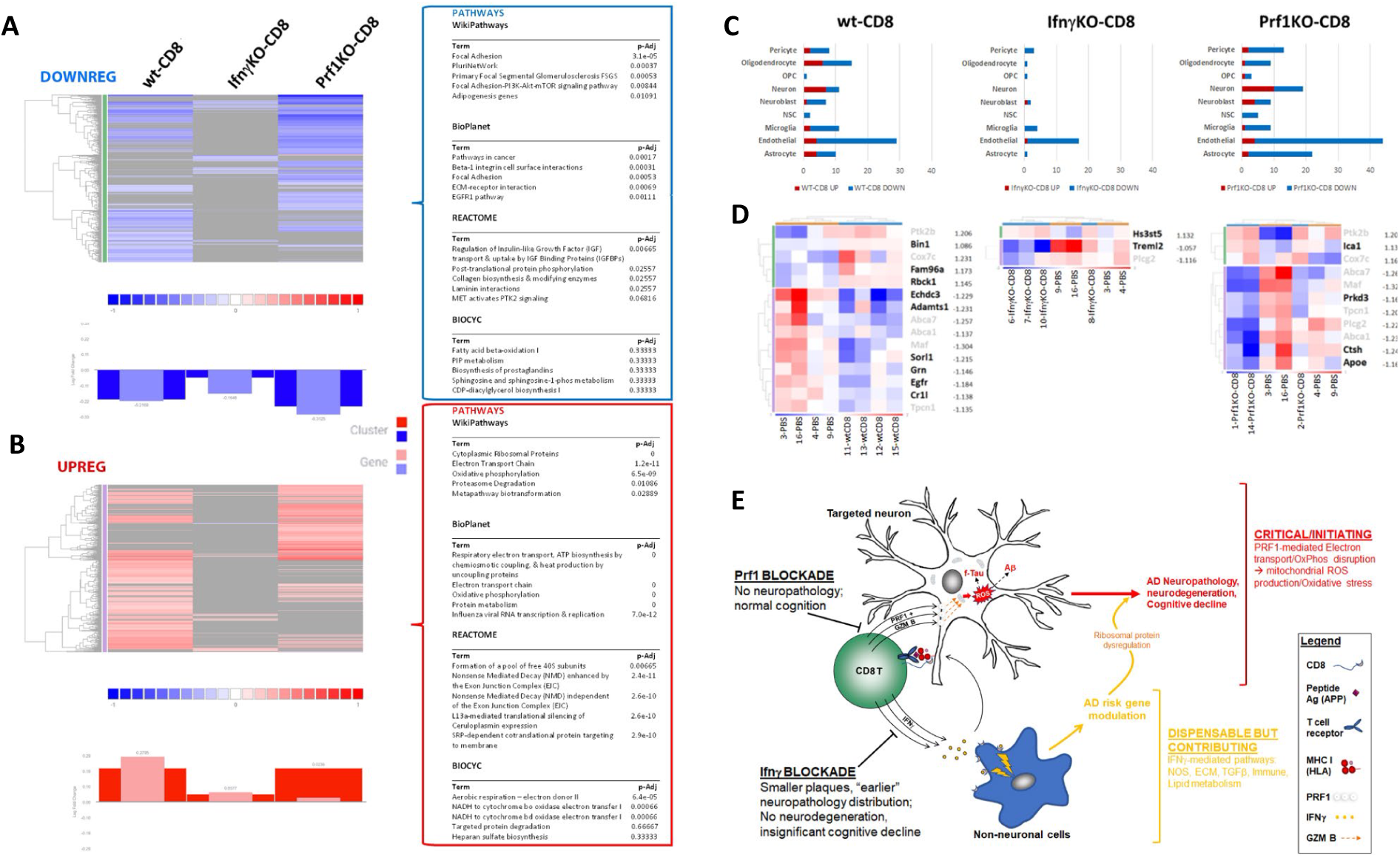
Gene expression changes in ^hi^T mouse brain. RNAseq analysis was performed on ^hi^T forebrains, yielding 1948, 309, and 1904 differentially expressed genes in wt-CD8, IfngKO-CD8 (all n = 4), and PrfKO-CD8 group (n = 3), respectively, relative to PBS. Upregulated (**a**) and downregulated (**b**) genes +/- 10% of PBS are depicted with associated cluster, gene, and ROSALIND knowledgebase analysis. Most cell type-specific genes were downregulated with the exception of neuron-specific genes, which were primarily upregulated (**c**). Differentially regulated genes among 84 AD-associated loci from GWAS studies (**d**). Mechanistic model of gene, pathway, and disease induction by ^hi^T cells based on combined pathological, knockout, and gene expression analysis (**e**).

### ^hi^T cell metrics in human Alzheimer’s disease

Given the varied impact of CD8 T cells on models of hereditary neurodegeneration, it was imperative that the potential impact of ^hi^T analogues was examined in human patients. We first examined levels of the larger “parental” memory/aging (KLRG1^+^) CD8 population in blood, and then the APP-specific KLRG1^+^ CD8 subpopulation derived from it, in HLA-A2^+^ individuals from 2 separate cohorts (Fig 5A, B-F). As expected, the parental CD8 population increased with age (Supplementary Materials fig S11) and also exhibited slight expansion in age-related cognitive decline or MCI-AD in both cohorts (Fig 5A, C; Supplementary Materials fig S12A,B). By contrast, APP-specific KLRG1^+^ CD8 T cells were markedly decreased in MCI and/or AD (Fig 5C, D), and all KLRG1^+^ T cell trends were maintained in both females and males (Supplementary Materials fig S13). The decrease in APP-specific KLRG1^+^ CD8 T cells correlated with poor cognitive performance in both cohorts independent of age (Fig 5E, F), exhibiting a linear correlation in the more sensitive MoCA test.

**Fig. 5:**
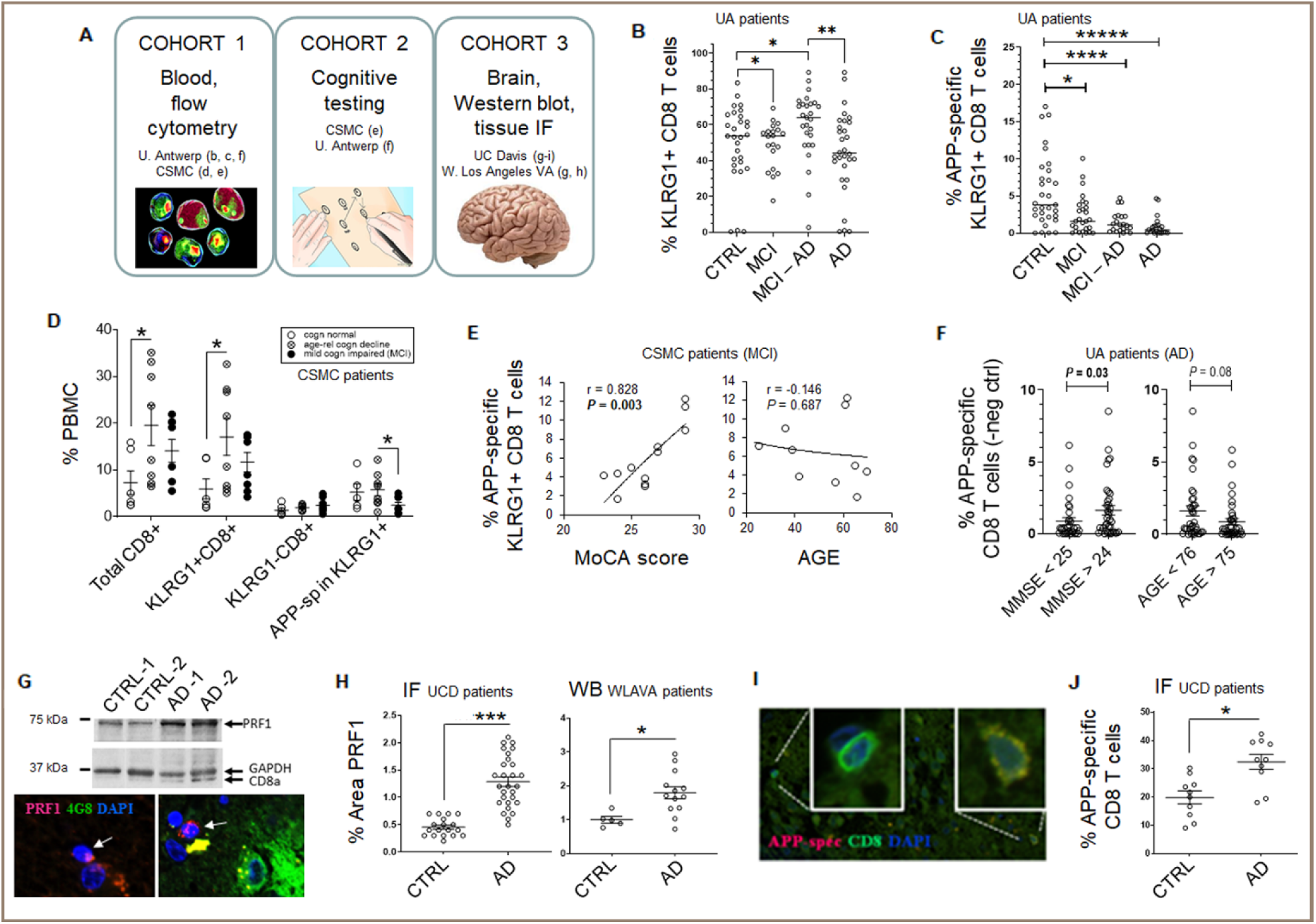
h^i^T parameters in human Alzheimer’s. Patient cohorts: University of Antwerp = “UA”; Cedars-Sinai Medical Center = “CSMC”; University of California, Davis = “UCD”; West Los Angeles Veteran’s Administration Hospital = “WLAVA” **(a)**. KLRG1^+^ (**b**) and APP_(471-479)_/HLA-A2-reactive KLRG1^+^ (**c**) CD8 T cells in CTRL, MCI ± CSF AD biomarkers (MCI, MCI–AD), and verified Alzheimer’s (AD) blood. T cell subpopulations vs. MoCA score (**d**), and correlation of APP_(471-479)_/HLA-A2-reactive KLRG1^+^ CD8 with cognitive test score and age (**e, f**). PRF1 Western blot (WB) and immunofluorescence (IF; **g)**, with quantifications in age-matched CTRL and AD brains. **(h**). APP_(471-479)_/HLA-A2-reactive CD8 staining (**i**) and quantification (**j**) in brain. Plots depict averages ± SEM. \**P* < 0.05, \*\**P* < 0.01, \*\*\**P* < 0.005, \*\*\*\**P* < 0.001 by 2-sided T-test, relative to CTRL unless otherwise indicated.

Cytolytic and antigen-specific T cell markers in AD brains were next examined within two independent patient cohorts (Fig 5A, G-J). Upregulation of Perforin1 in AD brain was evident in both patient cohorts, appeared coordinated with that of CD8 on Western blots, and exhibited the expected punctate cellular pattern by tissue immunofluorescence (Fig 5G, H). Similarly, T cells stained by CD8 and APP-specific pHLA multimers were more prevalent by tissue immunofluorescence in AD brain from the UCD cohort (containing both protein lysates and tissue sections), despite no pre-screening of specimens for HLA-A2 to which the APP-specific multimer binds (Fig 5I, J)). Taken together, these data suggest that parental CD8 T_RM_ in blood accumulate with aging, but house APP-specific CD8 T cells that are lost in proportion to cognitive decline in AD and prodromal conditions, just as they accumulate in AD brain.

Decreased APP-specific KLRG1^+^ CD8 T cells in blood correlated significantly with reduced Aβ1-42 and increased total Tau in CSF within the largest patient cohort (UA; Fig 6A), suggesting some correspondence with existing AD biomarkers. Accordingly, levels of these cells correlated uniquely with CSF Aβ1-42 exclusively in AD patients independent of age (Fig 6B). These trends suggested specificity for AD, and hinted at potential AD biomarker utility.

**Fig. 6:**
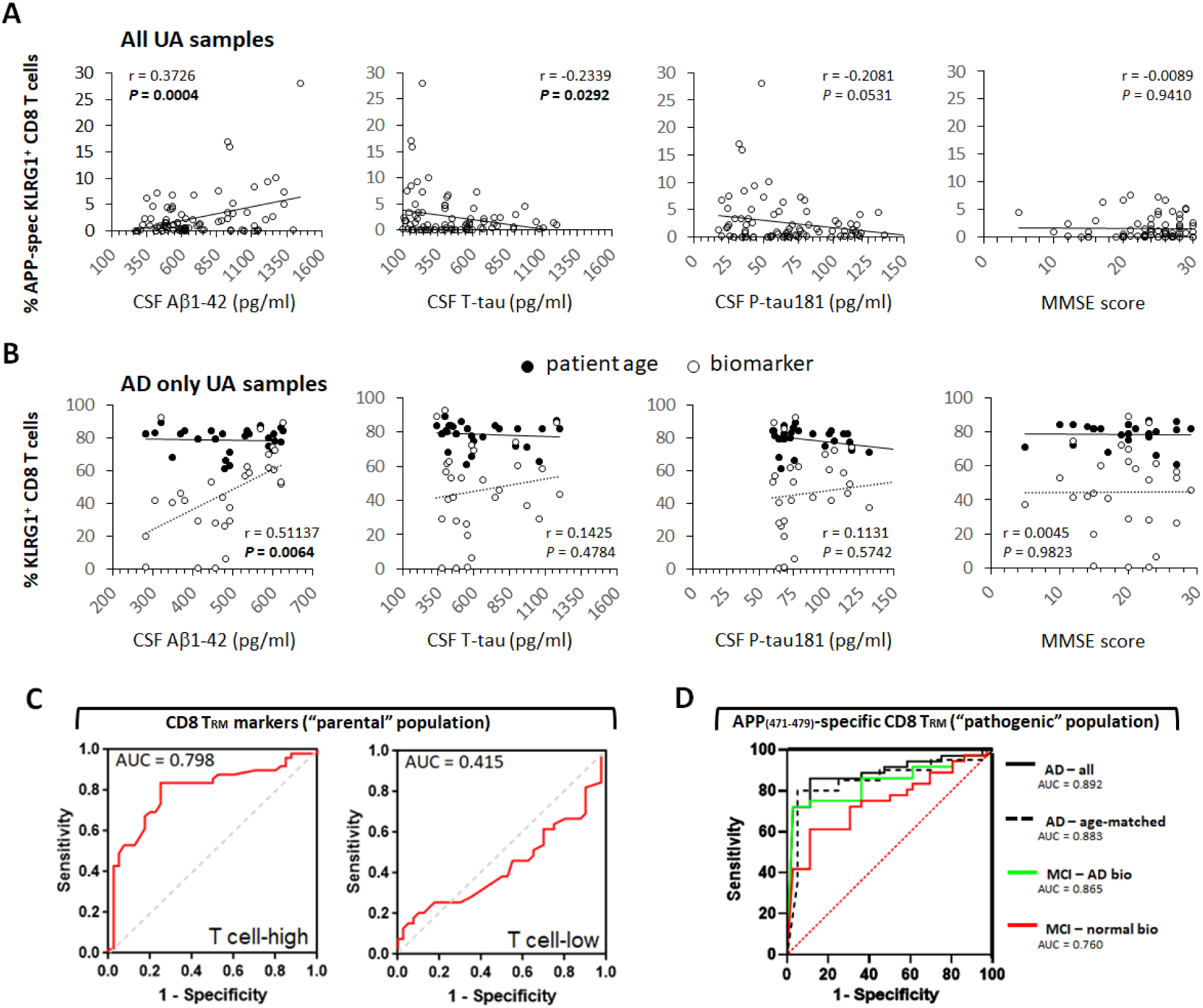
CD8 T_RM_ biomarker potential in human Alzheimer’s. Correlation of APP_(471-479)_/HLA-A2-reactive KLRG1^+^ CD8 levels with Aβ1-42, total-tau, and P-tau181 in CSF, and with MMSE score in all AU patients combined (**A**), and correlation of APP_(471-479)_/HLA-A2-reactive KLRG1^+^ % in CD8 levels or patient age with Aβ1-42, total-tau, and P-tau181 in CSF, and with MMSE score in AU Alzheimer’s patients only (**B**). KLRG^+^CD8^+^ T cell levels within lymphocytes exhibited a similar unique correspondence with CSF Aβ1-42, exclusively in AD patients (r = 0.6820, *P* < 0.00009). \**P* < 0.05, \*\**P* < 0.01, \*\*\**P* < 0.005, \*\*\*\**P* < 0.001 by 2-sided T-test with 1-sided T-test values depicted in parentheses, relative to CTRL unless otherwise indicated. Variance in CD103 (**C**). Combined variance of T_RM_ markers, CD8A, CD44, and CD103 yielded the indicated areas under the curve (AUC) in Receiver Operating Characteristic (ROC) plots of T cell-high and T cell-low samples from a publicly available patient dataset (**C**). AUC was not significantly altered when CD103 variance was used alone in this cohort (0.781, T cell-high; 0.548, T cell-low). ROC plots of APP_(471-479)_/HLA-A2 multimer-reactive KLRG1^+^ CD8 T cells in blood relative to normal aging controls, both from UA cohort (**D**). Mild Cognitive Impairment without (MCI-normal bio) and with (MCI-AD bio) CSF biomarkers consistent with AD, and confirmed AD patients ages 57-84 (AD-all). Area Under the Curve (AUC) is indicated. AD-age-matched indicates ROC analysis on 10 AD patients for whom precisely age-matched controls were available (+/- 1 year; n = 10). *P* < 0.001 for all curves except MCI – normal bio (*P* = 0.003). \**P* < 0.05, \*\**P* < 0.01, \*\*\**P* < 0.005, \*\*\*\**P* < 0.00005, 2-sided T-test.

To begin to assess such utility, we examined available data on blood from an additional independent patient cohort. Interrogating multiple CD8 T_RM_ markers (CD3D/CD3E/TCRz, CD8A/CD8B, CD103, CD122 or CD44), we verified significant upregulation at various stages of AD (Supplementary Materials fig 14A-C). By contrast, CD4 was inconsistently regulated in different cohorts. We separated the largest of these cohorts into T cell high and low groups based on median CD3 expression to exclude patients with age-related T cell lymphopenia, and those under 65 to exclude early onset AD patients and age-matched controls (Supplementary Materials fig 14C). Co-modulation of CD103, CD44, and CD8A as a group, as well as CD103 level alone, exhibited intriguing biomarker potential in Receiver Operating Characteristic (ROC) plots (Fig 6C). Moreover, APP-specific KLRG1^+^ CD8 T cell levels by flow cytometry exhibited markedly better biomarker potential than CD8 T_RM_ genes (Fig 6D), corresponding superiorly to MCI-AD than the highly touted plasma biomarker, pTau-217 (Supplementary Materials fig 15A-C), corresponded to AD itself (39). This demonstrates that the APP-specific CD8 T cell subpopulation and its parental KLRG1^+^ population are uniquely modulated in AD, and their levels can be uniquely useful as disease biomarkers.

## Discussion

The role of T cells in neurodegenerative disease has recently attracted considerable research attention. Nevertheless, many studies in Tg rodents have been contradictory to date, leaving questions of whether T cells dampen or exacerbate disease processes unresolved. Many of these studies use different transgenic mouse lines with distinct neuropathology, and none model AD induced by processes upstream of Aβ/pTau and the gene mutations promoting their deposition. Given that non-genetic AD encompasses 95-99% of patients, this is a critical shortcoming of current AD research.

The ^hi^T mouse model demonstrated several key features reminiscent of AD in humans. These include coupled accumulation of amyloid plaques and NFT-like fibrillar inclusions in tau^+^ neurons, as well as robust neuronal loss and profound cognitive decline, each demonstrated by at least three separate methodologies. These AD-like have not previously been induced in animal models by a single factor or transgene, as rodents typically fail to develop either NFT-like inclusions or robust neuronal loss without the addition of transgenes unrelated to human AD (1, 2, 40). Apart from these disease hallmarks, ^hi^T mice also exhibited finer features often seen in human AD, including cognitive loss progressing from hippocampus- to amygdala-dependent tasks over time **(26)**, progressive brain atrophy related to neuronal loss, unique association of cognitive loss with fibrillar tau pathology **(33)**, and modulation of multiple risk-associated genes.

Mechanistically, Perforin 1 in CD8 T cells was required for all aspects of neurodegenerative pathology, with the exception of most gene expression. While this potentially involved the target-lysis function of Perforin in T cells, we’d previously reported that Perforin also precluded the accumulation of T cells in brains of ^hi^T mice (19), a finding inconsistent with the minimal impact on gene expression its absence conferred on ^hi^T mouse brains. Follow up analysis showed that Prf1KO CD8 T cells did in fact accumulate and remain in ^hi^T brains long term, but at lower levels than wild-type CD8 T cells. While this offers an explanation for the changes in brain gene expression seen in Prf1KO-CD8 group mice, as well as pointing to a requirement for T cell Perforin in initiating AD-like pathophysiology, it does not discount the possibility that lower levels of brain T cells could be a contributing factor in the lack of neuropathology observed in Prf1KO-CD8 mice, particularly given that Perforin deficiency has been implicated in clinical disorders of T cell tissue homeostasis previously (41).

By contrast to Perforin 1, IFNγ in CD8 T cells appeared necessary to promote widespread neuronal loss and cognitive decline in ^hi^T mice, decreased the numbers and size of amyloid plaques, and limited their distribution and that of fibrillar tauopathy to brain regions impacted earlier in human AD. This parallels the impact of this cytokine of mouse models of amyloidosis (42). The apparent arrest of IfnγKO-CD8 group mice at a stage resembling early AD made them attractive to examine potential targets of T cell elimination in the absence of more widespread neuronal loss. CD8 T cells require MHC I on target cells to recognize and destroy them, but only small anatomically restricted subpopulations of neuronal lineage cells constitutively express MHC I without induction by cytokines such as IFNγ. Such cells include catecholaminergic neurons within locus coeruleus (35), among the earliest sites affected by AD neuropathology (43). Doublecortin (DCX)^+^ neuroblasts mediate adult neurogenesis, a process thought to be critically impaired early in AD, and also constitutively express MHC I (44-46). We hypothesized and then demonstrated that DCX^+^neuroblasts, as representatives of constitutively MHC I^+^ neuronal lineage cells in brain, were eliminated by CD8 T cells in the IfnγKO-CD8 group of ^hi^T mice. This suggests that neuronal lineage cells that constitutively express MHC I are initial targets of CD8 T cell elimination in the ^hi^T model, and that more widespread elimination of brain cells requires production of IFNγ in T cells, potentially inducing MHC I and conferring T cell sensitivity to ever widening groups of neural cells.

On a molecular level, Perforin1 is known to promote mitochondrial ROS and oxidative stress (very early events in AD) by facilitating Granzyme-mediated disruption of the electron transport chain in target cells (47, 48). Perforin 1 also promotes internalization of Aβ in neurons (49). By contrast, IFNγ is a well known pro-inflammatory effector, as well as a potent modulator of gene expression in cells expressing its receptor (36). The distinct involvement of these two genes is thus consistent with Perforin-mediated attack of neurons by ^hi^T cells, which simultaneously initiates limited neurodegeneration, mitochondrial oxidative stress, and inflammatory cytokine production in brain. IFNγ accelerates progression of the neuropathology through induction of MHC I and modulation of multiple risk-associated genes. CD4 T cells appear to play a dispensable role in induction and progression of neuropathology in ^hi^T mice, but appear to subtly influence the brain region damaged by ^hi^T_RM_, to the extent that it resembles a distinct AD behavioral subtype (50). As with the cellular mechanism outlined above, validation of this working molecular mechanism requires further detailed study.

While several aspects of the ^hi^T model paralleled features of human AD, it is not a perfect phenocopy of the human disease. For example, Aβ1−40 and diffuse plaques predominate in ^hi^T mice, whereas Aβ1−42 and compact plaques typically predominate in most forms of human AD. In addition, “ghost” tangles - remnants of neurofibrillary tangles in dead neurons - were not seen in ^hi^T mice. Both these differences may well be explained by ^hi^T mouse pathology occurring in the context of endogenous rather than transgenic human Aβ- and/or tau-encoding genes. Rodents, for example, are known to be deficient in clearing endogenous Aβ1-42, and exhibit Aβ1-40 preference in amyloid fibril formation **(51, 52)**, which could favor diffuse plaque and Aβ1-40 predominance in ^hi^T mice. Similarly, the absence of ghost tangles is very likely due to the virtual absence of the ghost tangle-promoting isoforms of MAPT, the gene encoding tau proteins, in adult mice **(53)**. In addition, the ^hi^T model was analyzed exclusively in female mice due to concerns over male-specific autoimmune dynamics in related strains (20, 21), making it imperative to test its relevance to human males and females with AD.

Reports of rodents exhibiting neurodegeneration linked to amyloid and fibrillar tau pathology have been published previously, most notably upon infection of mice with *Porphyromonas gingivalis* **(54)**. Introduction of multiple AD-associated human Aβ-related transgenes into rats (AD-Tg rats) has similarly been reported to produce neurodegeneration with NFT-like silver-stained neurons, although this model appears more relevant to dominantly inherited AD **(23)**. Evidence of NFTs in both these models, however, was extremely limited, reliant on single staining reagents, and was not statistically validated. For this reason, we sought to include one of these models as a control for NFT-like structures. In our hands, however, NFT-like structures failed to appear in the same AD-Tg rat model under staining procedures that readily revealed NFTs in ^hi^T mice and human AD brain alike. Further, neither infection nor transgenic models have offered mechanistic explanations for anatomically-specific disease etiology, generated pathophysiology corresponding to well-defined aspects of human AD progression, or demonstrate translational relevance more generally.

In addition to mirroring gross and fine aspects of human AD, ^hi^T mice successfully predicted that CD8 T_RM_-marker genes and Perforin protein are up-regulated in AD brain. Similarly, the model successfully predicted that APP-specific CD8 T cell levels would correspond to disease status in both females and males with AD and related MCI, as well as to cognitive decline in multiple cohorts. Finally, observations in ^hi^T mice led to the prediction that levels of ^hi^T cell analogues in blood would constitute uniquely useful tools for the clinical management of AD, with their utility for AD diagnosis particularly highlighted in this study. The role of T cells in neurodegenerative disease has recently attracted considerable research attention. Nevertheless, many studies in Tg rodents have been contradictory to date, leaving questions of whether T cells dampen or exacerbate disease processes unresolved. Many of these studies use different transgenic mouse lines with distinct neuropathology, and none model AD induced by processes upstream of Aβ/pTau and the gene mutations promoting their deposition. Given that non-genetic AD encompasses 95-99% of patients, this is a critical shortcoming of current AD research.

The ^hi^T mouse model is the first to replicate key aspects of AD via an inductive factor upstream of Aβ and other pathological hallmarks, and in addition exhibits clear translational relevance to sporadic AD. As such, it helps to dispel some of the confusion that surrounds the role of T cells on Alzheimer’s and related neurodegenerative diseases. Critically, the ^hi^T model reproduced compelling AD-like characteristics through introduction of age-related T cells and without the introduction of deterministic transgenes. Functional impairment of the T cells attenuated AD-like features and their analogues were uniquely associated with human AD. Thus, three critical criteria for causation (disease recapitulation by agent introduction into animals, disease mitigation upon agent impairment, and preferential association of agent and/or its dynamics to human disease) appear satisfied by this model. Nevertheless, the exclusive use of female nude mice qualifies such a conclusion and opens the possibility that additional factors associated with gender and/or strain genetics could also impact neurodegenerative induction. Similarly, human AD may in fact possess multiple distinct inductive events or processes. Hence, it will be critically important to discern the extent to which ^hi^T cells and related events contribute to AD induction and in what proportion of patients, if any, in future studies.

Conservatively, however, our study reveals the first evidence for the existence of a discrete and clinically useful factor upstream of Aβ that may contribute to amyloid- and tau-associated neurodegeneration. Further assessment of the potential of ^hi^T_RM_ metrics as diagnostic biomarkers for human AD nevertheless requires multi-center validation, prospective and longitudinal analyses that include more extensive non-AD dementia cohorts. It will be equally important to determine whether modulation of antigen-specific ^hi^T_RM_ presence or function can alter the course of AD-like neurodegeneration in therapeutic studies, and determine if effective treatments for AD critically modulate these cells in humans. Finally, continued examination of ^hi^T mice on multiple strain backgrounds, and harboring risk factors for AD or other age-related disorders, may lead to a more comprehensive understanding of age-related disease biology, pathology, and immune characteristics.

## Materials & Methods

### Animal subjects

Female C57BL/6, B6.Foxn1 mice, and congenic and/or syngeneic knockout strains (Jackson Labs) were housed in a pathogen–free vivarium under standard conditions on a 12-h light/12-h dark cycle with food and water *ad libitum.* Recipient animals were 8-10 week-old female B6.Foxn1, B6.Foxn1-AppKO, or B6.CD45.1-congenic mice; donors were 5-8 week-old females of the same strains. Specific numbers of animals used for all analytical methods are included in Supplemental Table 1. Cell derivation was randomized by pooling from ≥5 donors per experiment. Young (8-10 wk) and aged (15 months) male and female C57BL/6 and B6.CD103-knockout mice (n = 12 young; n = 7-8 aged) were used to study age-related cognitive decline. Donor, recipient, and unmanipulated animals were maintained in a pathogen-free facility under the Cedars-Sinai Department of Comparative Medicine, with all breeding and genetic screening conducted at Jackson Laboratories (Bar Harbor, ME).

### Adoptive transfer of T cells

Splenic CD8^+^ T cells from C57BL/6J female mice (5-7 weeks old) were purified using anti-CD8 or anti-CD4 immunobeads (Miltenyi Biotech, Sunnyvale, CA). 3 x 10^6^ purified CD8 or CD4 T cells, or 3 x 10^6^ native splenocytes for CD8+CD4 T cell hosts, were intravenously injected in 50 µl of PBS into female C57BL/6J or B6.Foxn1 nude hosts. Cell numbers were based on prior publications demonstrating homeostatic expansion at similar doses (19, 55, 56). Transfer efficiency into B6.Foxn1 hosts was validated by persistence of ≥5% CD8^+^, CD4^+^, or both CD8^+^ and CD4^+^ T cells, within splenic lymphocytes 3 weeks after injection (19). The order of treatments was randomized by alternating cell and control injections between individual recipients. For all subsequent analyses, performing investigators were blinded to both group definition and anticipated outcomes.

### Tissue processing

Brain and spleen were harvested from mice perfused with saline under deep anesthesia using a Ketamine and Xylazine (40-50 mg/kg i.p.) cocktail, until major organs such as liver and lungs lost color, and tissue was then excised for analysis. Upon removal of the whole brain from the cranium, the cerebellum, brainstem, and olfactory bulbs were removed, and remaining brain tissue was weighed on a Mettler balance for standardized brain mass assessment. Brains were sectioned 1 mm to the right of the longitudinal fissure (midline). Right hemispheres were flash frozen in -80°C conditions for protein studies, followed by homogenization in Cell Lysis Buffer (Cell Signaling Technologies, MA), and centrifugation of nuclei. Cell lysates were separated into Triton-soluble, Sarkosyl-soluble and Sarkosyl-insoluble fractions using sequential incubations with 10% (wt/V) salt sucrose solution and 1% (wt/v) sarkosyl Salt Sucrose Solution. Left hemispheres were immersion fixed in 4% paraformaldehyde (duration?) and reserved for immunohistochemical staining.

### Antibodies for tissue staining and western blot analyses

Free-floating brain sections (8-14 µm thick) were mounted onto slides and blocked with Protein Block (Serum-Free, Dako, CA) for 1h at RT. Sections were incubated at 4°C overnight with primary antibody in Protein Block (Dako, CA). Sections were rinsed 4x in PBS, and incubated 90 min in fluorochrome- or biotin-conjugated secondary antibody, with or without curcumin (0.01% in PBS), or with ThioS alone (1% in PBS). Sections were washed, coverslipped, and mounted with ProLongGold anti-fade media with DAPI (Invitrogen). Bright-field and fluorescent images were obtained using a Zeiss AxioImagerZ1 with CCD camera (Carl Zeiss Micro imaging). Image analysis of micrographs was performed with ImageJ (NIH). Anti-Aβ/APP antibody (ab14220, Abcam for 3-week time point; clone 4G8, Chemicon for all others) was used at 1:500 for immunohistochemistry (IHC) and 1:1000 for Western blot (WB). Anti-pTau pS199/202 antibody (Invitrogen) was used at 1:50 for IHC and 1:100 for WB, with PHFs confirmed with Phospho-PHF-tau pSer202+Thr205 Antibody (AT8), used at 1:2000 in WB. AT8 is extensively published as pTau-reactive in mice and humans, with the caveat that it can react with endogenous mouse immunoglobulins (Ig; IgG H chain) to yield non-specific bands around 50kDa – larger Ig bands are not seen in reducing SDS/PAGE gels and derivative blots, and non-tau AT8 bands disappear when endogenous Ig is adsorbed (57). We thus used reducing SDS/PAGE gels only, and examined species between 75-100 kDa with AT8 in Western blots, eliminating the possibility of non-specific bands reactive to mouse Ig. The Invitrogen anti-Tau antibody was used to examine lower MW pTau species, to exclude issues in this size range associated with AT8. Due to marker size, pTau WB signal was normalized to that of β-actin (clone AC-74, Sigma), with GAPDH used for normalization of all other markers. Anti-GFAP (Dako) was used at 1:250 for IHC and WB. Anti-NeuN antibody (Chemicon) was used at 1:100 for IHC and WB. Anti-Iba1 (Wako, Ltd.) was used at 1:200 for IHC. Anti-CD8 (clone 53-6.72, BD Pharmingen) was used at1:100 for IHC and 1:1000 for WB. Anti-doublecortin (DCX; polyclonal sc-8066, Santa Cruz Biotechnology) was used at 1:2000 for immunofluorescence). All secondary antibodies (HRP, Alexa Flour-488, -594, -647; Invitrogen) were used at 1:200 for IHC and 1:2000 for WB. Multimer generation & use: dextramers of established epitopes for self/brain antigen (Trp-2-DCT_(180-188)_/H-2K^b^), and/or custom APP epitopes with predicted affinities < 100nM (NetMHC version 3.4; APP_(470-478)_/H- 2D^b^), were manufactured by Immudex.

### Western blot for amyloid, tau, neural and immunological markers

Triton-soluble cell lysates were electrophoretically separated on 12% Tris-HCl Precast Gels (Bio-Rad), and blotted onto 0.2 µm nitrocellulose membranes. Membranes were blocked with bovine serum albumin (BSA), incubated in sequential primary and secondary antibody dilutions for 1 hr at room temperature with ≥3 washes, developed with enhanced chemiluminescence substrate (GE Healthcare Biosciences; Pittsburgh, PA), and exposed onto Amersham Hyperfilm (GE Healthcare Biosciences; Pittsburgh, PA).

### ELISA

Supernatant from homogenized brain tissues was used for Triton-soluble Aβ. Insoluble pellets from Triton-homogenized brain were resuspended in 10 volumes 5M Guanidine HCl 4 hr to generate Guanidine-soluble Aβ. Triton- and Guanidine-soluble samples were subjected to analysis using species- and isoform-specific antibodies in Soluble and Insoluble Aβ ELISA (Invitrogen, Life Technologies; Grand Island, NY; mouse-specific Aβ40 ELISA Catalog # KHB3481; mouse-specific Aβ42 ELISA Catalog # **KMB3441**). Absorbance was read on a SPECTRAmax Plus384 microplate reader (Molecular Devices, Sunnyvale, CA) with data analyzed in Graphpad PRISM (Graphpad Software; San Diego, CA).

### Flow cytometry

Purified T cells stained with respective Abs were analyzed by three-color flow cytometry (FACScan II; BD Biosciences, San Jose, CA) to assess purity. Antibodies were incubated with whole-spleen single-cell suspension in PBS with 5% FBS, on ice for 30 min, followed by a wash with the PBS with 5% FBS. Subsequently, 100,000–300,000 flow events were acquired.

### Gallyas silver staining

Gallyas silver stain was used to visualize fibrillar aggregates. Free floating brain sections were placed in 5% Periodic Acid for 3 min, washed twice and placed in Silver Iodide solution 1 min, followed by incubation in 0.5% Acetic Acid 5 min (2X), and rinsing with dH_2_0. Sections were incubated in developer for ∼10 min until sections were pale brown/gray, and stopped in 0.5% acetic acid for 5 min, rinsed in dH_2_O and mounted. Stained sections were examined by microscopy. Stained neurons were counted from CA2 of hippocampus, and their proportions within total neurons visually quantified in triplicate from entorhinal and cingulate cortex.

### Neuronal counts

Whole-number neuronal estimates were performed using the optical fractionator method (58) with stereological software (Stereo Investigator; MBF Bioscience). Para-median sagittal serial sections spaced 50 µm apart were stained with NeuN. CA1, CA2, CA3 and other regions of interest were defined according to the Paxinos and Watson mouse brain atlas. A grid was placed randomly over the ROI, and cells were counted within three-dimensional optical dissectors (50 µm 50 µm 10 µm) using a 100x objective. Within each dissector, 1 µm guard zones at the top and bottom of section surface were excluded. Estimated totals weighted by section thickness were obtained with Stereo Investigator software, yielding a coefficient of error 0.10.

#### Behavioral testing - general

Open Field testing was performed preceding all other behavioral tests, at 3, 6, and 13 months post-cell or -control injection. Testing order was randomized by alternating control and treatment group animal runs. Testing started at the same time (+/- 1.5 hr) for tests run on more than one day, with early and late times alternated for inter-group randomization.

#### Open field test

Testing was carried out in an Open Field apparatus made up of an open topped, clear Plexiglas box, measuring 40.64 cm x 40.64 cm and 38.1 cm high. Two rings of photobeams and optical sensors surrounded the box. The optical sensors were connected to a computer by way of an input matrix. Each mouse was placed into the box, and beam interruptions were automatically recorded as a measure of locomotor activity. Each mouse was tested in the box for a period of 30 min.

#### Barnes maze test

Barnes Maze (BM) testing was performed a single time only, 14 months post-cell or –control injection. The BM test is a hippocampus-dependent, spatial-learning task that allows subjects to use spatial cues to locate a means of escape from a mildly aversive environment (i.e. the mice are required to use spatial cues to find an escape location). Mice were assessed for their ability to learn the location of an escape box over the course of 9 days in the BM apparatus (31, 59). The escape hole is constant for each mouse over the five training days. Each mouse was tested three times per day (3 trials) for 4 days, followed by no testing for 2 days, and re-testing on day 7. A 35-60 min inter-trial interval separates each trial. Each trial began by placing one mouse inside a start box with a bottomless cube positioned centrally on the maze. After 30 seconds, the start box was lifted and the mouse was released from the start box to find the one hole with access to the escape box. Two fluorescent lights located approximately 4 feet above illuminated the testing room. Each trial lasted up to 4 min or until the mouse entered the escape box. The experimenter guided mice that failed to find the escape hole within 4 min, to the correct hole after each training test. Once the mouse entered the escape box, it was allowed to remain in the box for 1 min. Following the 7^th^ day of testing, and never on the same day, mice were tested an additional two-days, in which the escape box was placed in the reverse position on day 8, and replaced in the original position on day 9. The same exact testing procedure was applied to all mice in all groups. The maze and all compartments were cleaned thoroughly with isopropyl alcohol to remove any olfactory cues after each trial, and prior to each day of testing. Additional randomization of alternating escape compartment location between each animal per group, and between each of 3 daily training tests per animal, was employed for this test.

#### Y-maze spontaneous alternation behavior

Mice were tested for SA a single time only, at 12 months post-cell or –control injection. Y-Maze Spontaneous Alternation Behaviour (SAB) is used to assess working memory. SAB was measured by individually placing animals in one arm of a symmetric Y-maze made of opaque black acrylic plastic (arms: 40 cm long, 4 cm wide; walls: 30 cm tall), and the sequence of arm entries and total number of entries recorded over a period of 8 min.

#### Flinch-jump/fear conditioning tests

Flinch-jump/Fear Conditioning freezing times were determined 6 and 11 months post-cell or – control injection. We first determined there were no significant differences in the nociceptive threshold (pain sensitivity) across treatment groups using the Flinch-Jump Test. Pavlovian Fear Conditioning was then used to assess learning and memory regarding aversive events. The apparatus (Freeze Monitor™, San Diego Instruments, San Diego, CA) consisted of a Plexiglas box (25.4 x 25.4 x 31.75 cm high) with a stainless-steel grid floor. An acoustic stimulus unit is located on top of the box, and the box is ringed with photo beams and optical sensors. The optical sensors were connected to a computer by way of an input matrix, and the number of beam interruptions is automatically recorded. For testing, on day 1 individual mice were placed into the test box, and allowed to habituate for 3 min. At 3 min a tone was presented for 30 sec. Then, 30 sec after termination of the tone, a 0.5 sec foot shock (intensity = mean jump threshold for the treatment group determined by the Flinch-Jump Test) was delivered. The mouse was then removed from the box and returned to its home cage for 2 min. The chamber was cleaned and the animal returned to the chamber where the procedure was repeated. The freeze monitor apparatus recorded freezing times throughout the procedure (absence of movement for 5+ seconds, resulting in no beam breaks). On day 2, context retrieval was determined by placing the mouse into the same test box where it previously received a tone and foot shocks, but here the tone and foot shocks were not presented. Freezing time was measured over a 10-min period. On day 3, cue conditioning was measured after inserting a triangular, plexiglass box into the test box. The mouse was placed into the triangular chamber where they had not previously received tone or foot shocks, but after 1 min the auditory tone was delivered for 30 sec and freezing time measured for 10 min.

### RNAseq and gene expression analysis

rRNA was depleted with NEBNext® rRNA Depletion Kit v2 and libraries were prepared with NEBNext® Ultra™ II Directional RNA Library Prep Kit. Data was analyzed by ROSALIND® (https://rosalind.bio/), with a HyperScale architecture developed by ROSALIND, Inc. (San Diego, CA). Reads were trimmed using cutadapt. Quality scores were assessed using FastQC. Reads were aligned to the Mus musculus genome build mm10 using STAR. Individual sample reads were quantified using HTseq and normalized via Relative Log Expression (RLE) using DESeq2 R library. Read Distribution percentages, violin plots, identity heatmaps, and sample MDS plots were generated as part of the QC step using RSeQC. DEseq2 was also used to calculate fold changes and p-values and perform optional covariate correction. Clustering of genes for the final heatmap of differentially expressed genes was done using the PAM (Partitioning Around Medoids) method using the fpc R library. Hypergeometric distribution was used to analyze the enrichment of pathways, gene ontology, domain structure, and other ontologies. The topGO R library, was used to determine local similarities and dependencies between GO terms in order to perform Elim pruning correction. Several database sources were referenced for enrichment analysis, including Interpro, NCBI, MSigDB, REACTOME, WikiPathways. Enrichment was calculated relative to a set of background genes relevant for the experiment. Sequencing data are deposited at https://jumpgate.caltech.edu/runfolders/volvox02/nextseq/220627_VH00472_82_AAC33CJM5/Analysis/1/Data/fastq/. Passcode access to sequencing data will be provided on request.

### Human Subjects

Cohort 1: 40 control individuals (CTRL; normal aging – normal CSF profile; 16M, 24F; avg age 60.2 yr [49-88]); 52 MCI patients with an AD-characteristic CSF biomarker profile (MCI-AD; 20M, 32F; avg age 75.5 yr [49-88]); 36 MCI patients not displaying an AD-characteristic CSF biomarker profile (MCI; 17M, 19F; avg age 62.22 yr [42-82]); 50 sporadic AD patients with an AD-characteristic CSF biomarker profile (AD; 26M, 24F; avg age 76.08 yr [57-88]). AD biomarker positivity (in-house validated cut-off values of biomarkers were applied: Aβ_1-42_ <638.5 pg/mL, T-tau>296.5 pg/mL, P-tau_(181P)_ >56.5 pg/mL) was determined by means of commercially available single-analyte ELISA kits (INNOTEST® β-AMYLOID (1-42), INNOTEST® hTAU-Ag, and INNOTEST® PHOSPHO-TAU (181P); Fujirebio Europe). CSF samples were collected at Middelheim General Hospital (Antwerp, Belgium) and centers referring to the Neurobiobank of the Institute Born-Bunge (NBB-IBB; n° BB190113) according to standard collection protocols as described previously (60). CSF was obtained by lumbar puncture (LP) at the L3/L4 or L4/L5 interspace. CSF samples were collected in polypropylene vials (Nalgene cat.no.5000–1020 (1.5 mL) and 5000–0050 (4.5 mL)), immediately frozen in liquid nitrogen, and subsequently stored at –80°C until analysis. In addition, blood samples following venepuncture were collected after LP, of which 2-3 serum, 3-4 plasma and 2-3 total blood aliquots were stored (1.5mL for each aliquot).

MCI patients underwent LP at baseline as part of their diagnostic work-up. The inclusion criteria for the control group were: [1] no neurological or psychiatric antecedents and [2] no organic disease involving the central nervous system following extensive clinical examination. MCI patients were diagnosed applying Petersen’s diagnostic criteria (61), i.e., [1] cognitive complaint, preferably corroborated by an informant; [2] objective cognitive impairment, quantified as performance of more than 1.5 SD below the appropriate mean on the neuropsychological subtests; [3] largely normal general cognitive functioning; [4] essentially intact activities of daily living (basic and instrumental activities of daily living were determined by a clinical interview with the patient and an informant); and [5] not demented (62). AD dementia was clinically diagnosed according to the NINCDS/ADRDA and IWG-2 criteria (63) APOE allele status was 40% 3/4, 40% 4/4, 10% 3/3, 10% 2/4 for AD group (10 tested/50 total); 55% 3/4, 20% 4/4, 15% 3/3, 10% 2/4, 0% 2/3 for MCI-AD group (20 tested/52 total); 31% 3/4, 0% 4/4, 46% 3/3, 8% 2/4, 15% 2/3 for MCI-normal group (13 tested/36 total); 22% 3/4, 0% 4/4, 68% 3/3, 0% 2/4, 8% 2/3 for control group (37 tested/40 total). Mean MMSE score was 20 (5-29) +/- 5 for AD group (48/50 tested); 24 +/- 3 (15-30) for MCI-AD group (51/52 tested); 26 +/- 3 (18-30) for MCI-normal group (33/36 tested); 28 +/- 4 (17-30) for control group (9/37 tested). All included subjects were of Caucasian ethnicity.

Cohort 2: Cognitive Testing. 29 self-referred memory clinic patients (average age 49; 22-70) from Cedars-Sinai Dept. of Neurosurgery were clinically diagnosed as normal (n = 6), MCI (n = 18), dementia (n = 3), or uncertain (n = 2; diagnosis established solely on basis of cognitive testing), and followed up with cognitive (Montreal Cognitive Assessment; MoCA) testing (n = 22). Cognitively normal = MoCA score 29-30; age-related cognitive decline = MoCA score 26-28; MCI = MoCA score < 26. HLA-A2-negative patients were determined by flow cytometry and excluded from analysis, as were patients with ≤2.5% CD8^+^ cells in lymphocyte gates.

Cohort 3: Brain Western and IHC. Hippocampal lysates from 13 autopsy-confirmed Braak stage IV (n = 4) and Braak stage V-VI (n = 9) sporadic AD patients, and 5 age-matched normal controls were run on Western blots using anti-CD8 or anti-PRF1. Hippocampal sections from 10 autopsy-confirmed Braak stage V-VI sporadic AD patients, and 10 age-matched normal controls were stained with anti-CD8-flourescein plus APP_(471-479)_/HLA-A2-PE. HLA-A2-negative samples were not excluded from Western and IHC/IF analysis.

### Statistical analysis

Quantification and stereological counting procedure for cell numbers or area (µm^2^) of Aβ plaque, GFAP^+^, Iba1^+^ or Perforin1^+^ cells were analyzed in six to eight coronal sections from each individual, at 150-µm intervals (unless otherwise indicated), covering 900–1200 µm of the hippocampal and cortical areas. Specific fluorescence signal was captured with the same exposure time for each image and optical sections from each field of the specimen were imported into NIH Image J and analyzed as above. GraphPad Prism (version 5.0b; San Diego, CA, USA) was used to analyze the data using ANOVA and T-Tests with Welch’s correction (no assumption of equal variance). In all histograms, average ± SEM is depicted.

Data from Open Field and Barnes Maze tests were analysed by 2-sided T-Test for individual test points, and by ANOVA on test curves, when normal distribution/*P* > 0.05 of data was verified in Anderson-Darling, D’Agostino & Pearson, Shapiro-Wilk, and/or Kolmogorov-Smirnov tests. 1-sided T-Test was used to analyze Y-maze/SAB. Mann Whitney test was substituted for T-Tests when non-normal distribution/*P* < 0.05 was indicated in Anderson-Darling, D’Agostino & Pearson, Shapiro-Wilk, and/or Kolmogorov-Smirnov tests. Flinch-Jump and Fear Conditioning Tests was normalized, first within each group to the average of the initial two tests in training on day 1, and then within all experimental groups to the average contextual or cue values of PBS controls, expressed as percent of control, and analyzed by ANOVA, followed where appropriate by Newman-Keuls tests to detect differences among treatment groups.

Sample sizes for PrfKO-CD8 and IfnγKO-CD8 groups were calculated *a priori* for each metric using means and standard deviations of PBS and wt-CD8 groups for anticipated effect sizes, with alpha 0.05, and >95 confidence. Calculated n plus ≥1 were then used for PrfKO-CD8 and IfnγKO-CD8 groups.

Pre-determined exclusions included sections or samples with no discernible background signal, and values within each group ≥2 standard deviations above or below the median/group. Subject numbers and methods of reagent validation are shown in Table S1.

### Study approval

All animal procedures were approved prior to performance by the Cedars-Sinai Institutional Animal Care and Use Committee. The Cedars-Sinai Institutional Review Board designated the analysis of de-identified human brain specimens from UC Davis exempt from committee review. Brain specimens were collected, stored, and disseminated with prior approval by the UC Davis Medical Center Institutional Review Board. Sampling for cohort 1 was approved by the Medical Ethics Committee of the Hospital Network Antwerp (ZNA), Antwerp, Belgium (approval number 2805 and number 2806).

## Acknowledgments & Funding Sources

We gratefully acknowledge support in conducting behavioral tests from the Cedars-Sinai Research Institute Biobehavioral Core, Igor Antoshechkin for RNAseq performance, Jeremy Sanders and Sol Katzman for guidance in gene expression analysis, and Ms. Hannah Schubloom and Mia Oviatt for excellent administrative support and editing.

## Funding

National Institutes of Health grant (UC Davis Alzheimer’s Disease Center) P30AG10129 (L-WJ)

National Institutes of Health grant R21NSO54162 (CJW)

National Institutes of Health grant R21AG033394 (RMC)

Cedars-Sinai Medical Center Biobehavioral Core (RNP)

Joseph Drown Foundation (CJW)

Maxine Dunitz Neurosurgical Institute (CJW)

Maxine Dunitz Neurosurgical Institute (DKI)

## Author contributions

Conceptualization: RMC, KLB, DKI, BAW, CJW

Methodology: AP, AR, MJ, RC, NG, GD, AM, DG, HS, DVD, YV, HDR, DKI, BAW, CJW

Investigation: RMC, RNP, L-WJ, JAR, DKI, BAW, CJW

Visualization: AR, NG, DKI, CJW

Funding acquisition: RMC, RNP, DKI, KLB, CJW

Project administration: CJW

Supervision: KLB, PPDD, DKI, CJW

Writing – original draft: AP, AR, CJW

Writing – review & editing: RMC, JAR, CJW

## Data and materials availability

Results and raw data will be made available upon request. Model Organisms and/or the means to generate them will be made generally available for research (non-commercial) use.

