## Supplementary Information for "Antigen-specific age-related memory CD8 T cells induce and track Alzheimer’s-like neurodegeneration"

BioRxiv

**The PDF file includes:**

Figs. S1 to S16

Table S1

**A**

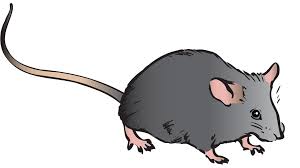

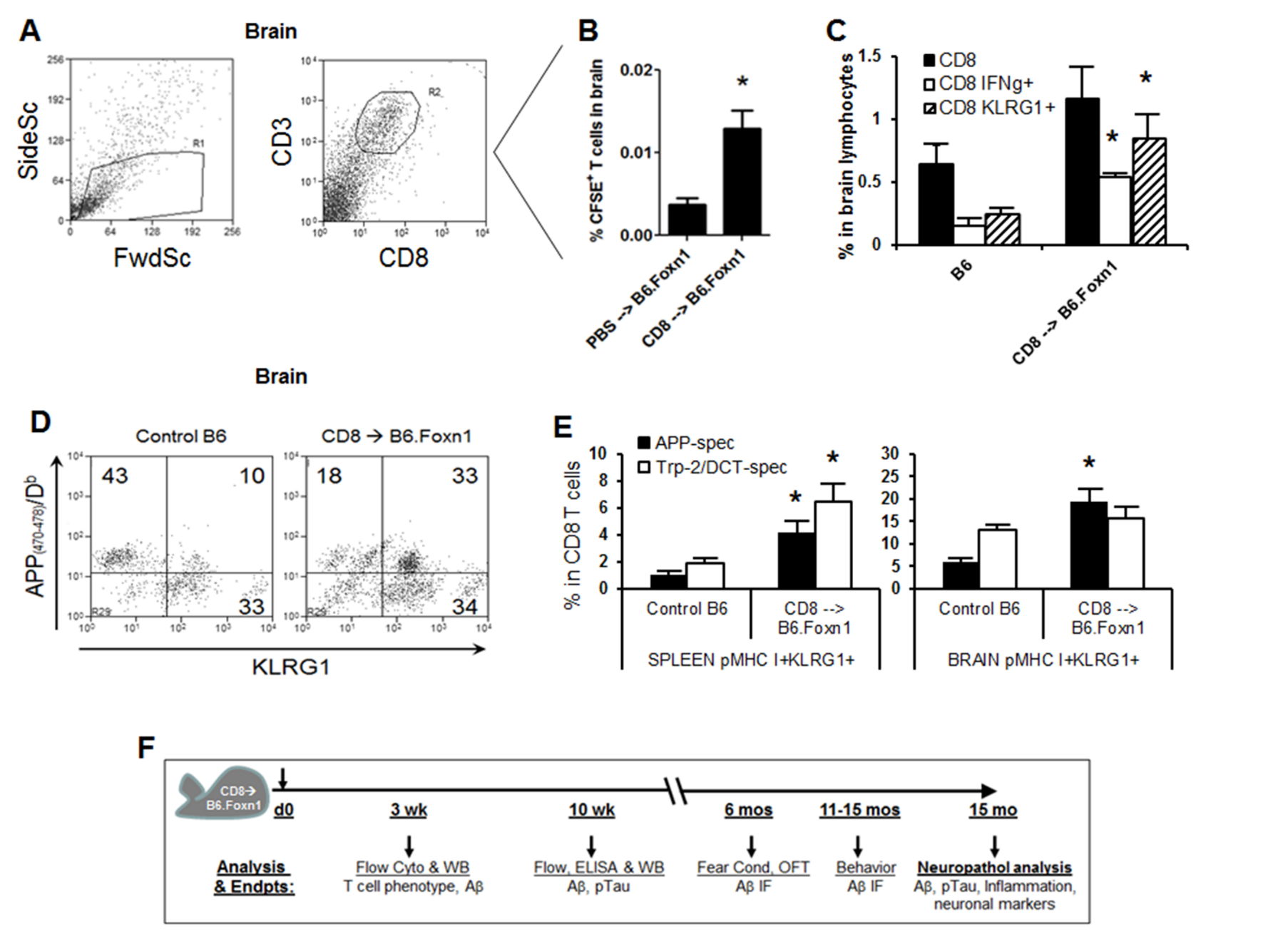

**B**

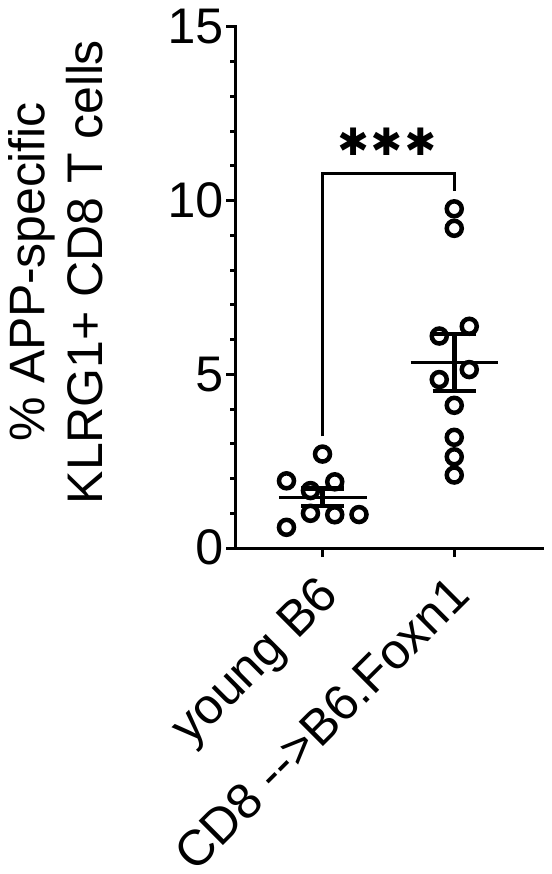

Fig. S1: Flow cytometry of APP-specific CD8 T cells in mice and humans. Analysis time line in CD8

T cell-injected B6.Foxn1 recipients (A). Levels of mouse APP_(470-478)_/H-2D^b^ and human APP_(471-_

_479)_/HLA-A2 were quantified within KLRG1^+^ CD8 T cells by flow cytometry in mouse and human

blood, respectively (B). Young B6 = APP-reactive T cells within splenic lymphocytes from 8-10 week-

old C57BL/6 females; CD8 🡪 B6.Foxn1 = APP-reactive T cells within splenic lymphocytes from 10-

12 week-old B6.Foxn1 female recipients of purified CD8 T cells from 6-8 week-old C57BL/6 females

3-5 weeks prior (****P* < 0.005 in 2-side Welch’s T-test).

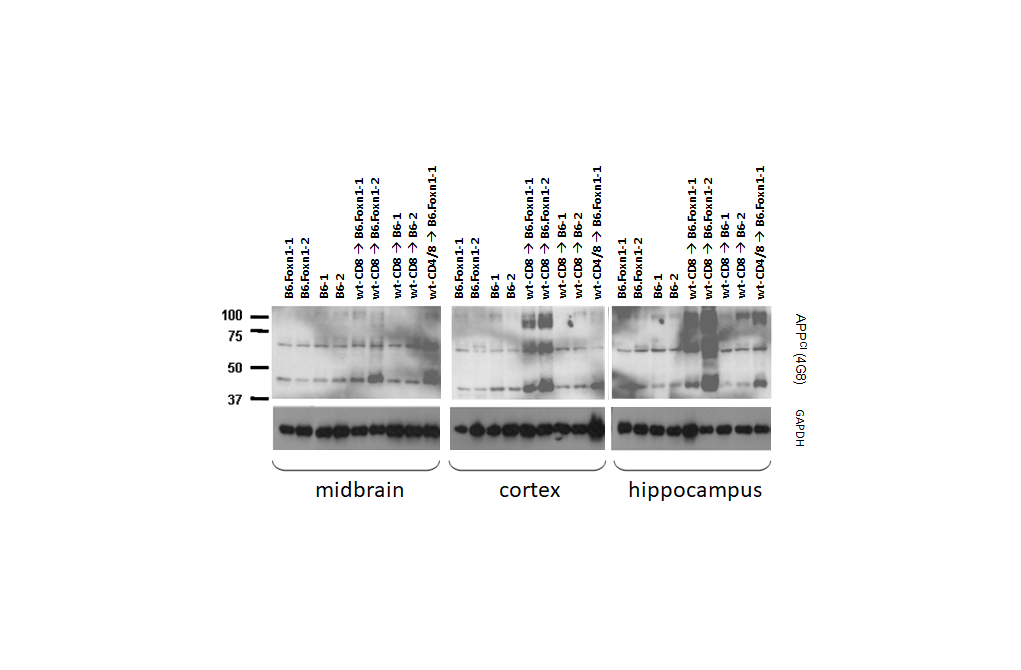

**D**

**A**

**B**

**C**

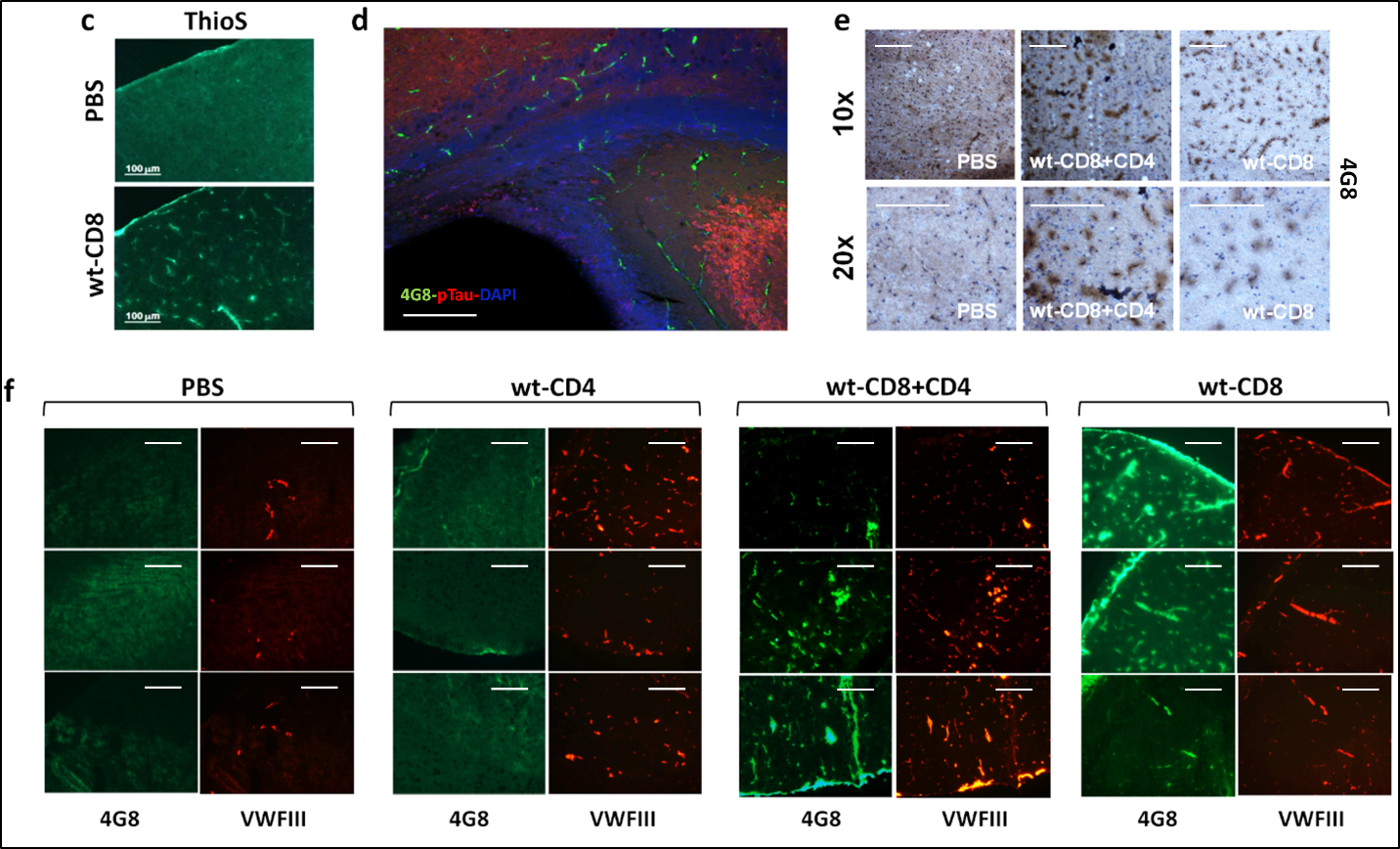

**E**

**Fig. S2: Analysis scheme, APP cleavage products, and vascular Aβ in brain after T cell injection.** Western analysis of APP cleavage products/Aβ oligomers with 4G8 antibody in dissected regions of mouse brain 10 weeks after injection with 3 x 10^6^ purified T cells (**A**). CD4/8 indicates injection of ~85% CD4 and 15% CD8, with peripheral proportions verified by flow cytometry. Immunofluorescence staining for ThioS in B6.Foxn1 cortex 6 months post-injection, exhibiting staining reminiscent of vasculature in wt-CD8 but not PBS group mice (**B**). Immunofluorescence staining for 4G8, phospho-tau, and DAPI in wt-CD8 group periventricular/hippocampal region 6 months post-injection, exhibiting vascular staining pattern (**C**). Immunohistochemical staining showing positive cortical 4G8 staining in wt-CD8+CD4 and wt-CD8 groups, but not PBS group, 6 months post-injection (**D**). Aβ (4G8) and Von Willebrand Factor III (VWFIII) staining in B6.Foxn1 cortex 6 months after injection, confirming deposits of aggregated vascular Aβ in nude mice harboring ^hi^T cells (**E**). All measurement bars are 100 μm in length.

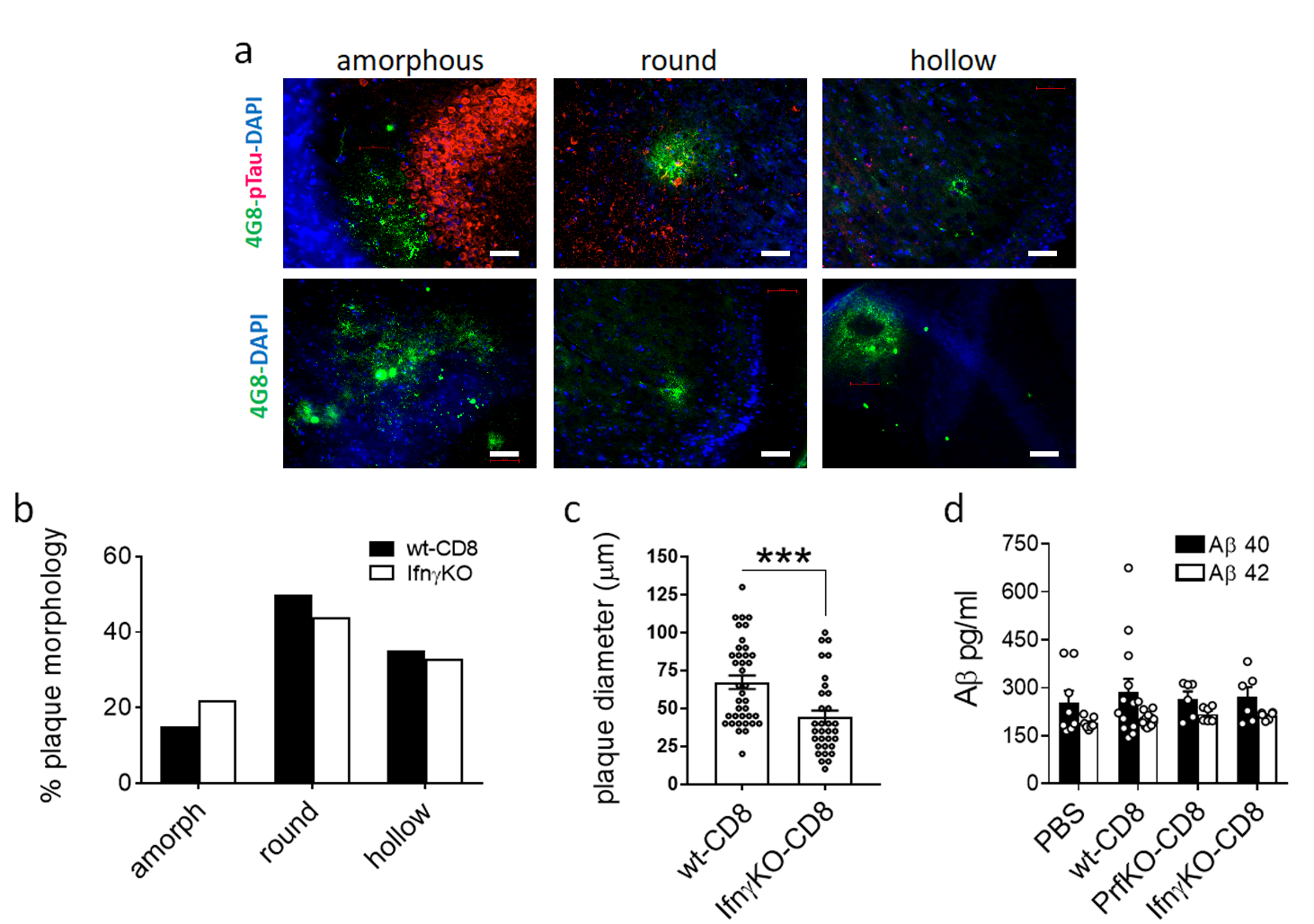

**Fig. S3: Aβ accumulation in nude mouse brain after CD8 T cell injection.** Forebrain ELISA of GuanidineHCl-soluble Aβ in B6.Foxn1 brain 15 months after CD8 T cell or control injections.

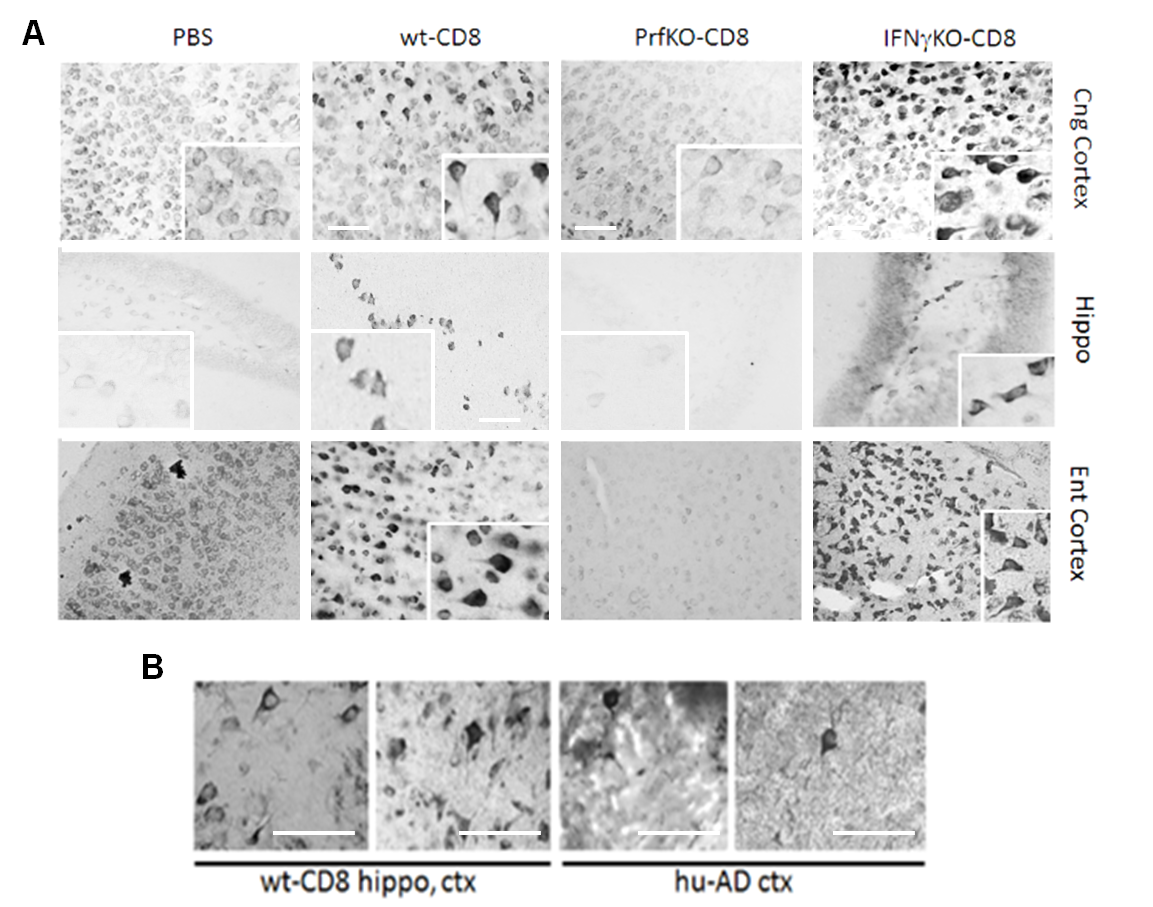

**Fig. S4: Silver stained neuronal structures in experimental groups.** Gallyas silver staining of cortical and hippocampal brain regions, showing typical neurofibrillary tangle (NFT) morphology in wt-CD8 and IFNγKO-CD8 group mice (insets) 15 months after cell injection. Background silver staining was occasionally evident in PrfKO-CD8 or PBS group mice, but did not exhibit similar NFT morphology (insets). Individual images were derived from different mice within each group (**A**). Comparison of Gallyas^+^ structures in nude mice harboring ^hi^T cells (wt-CD8) hippocampus (left) and cortex (ctx, right), to those in cortex of human severe AD (Braak stage VI; **B**). Measurement bars are all 50μm in length.

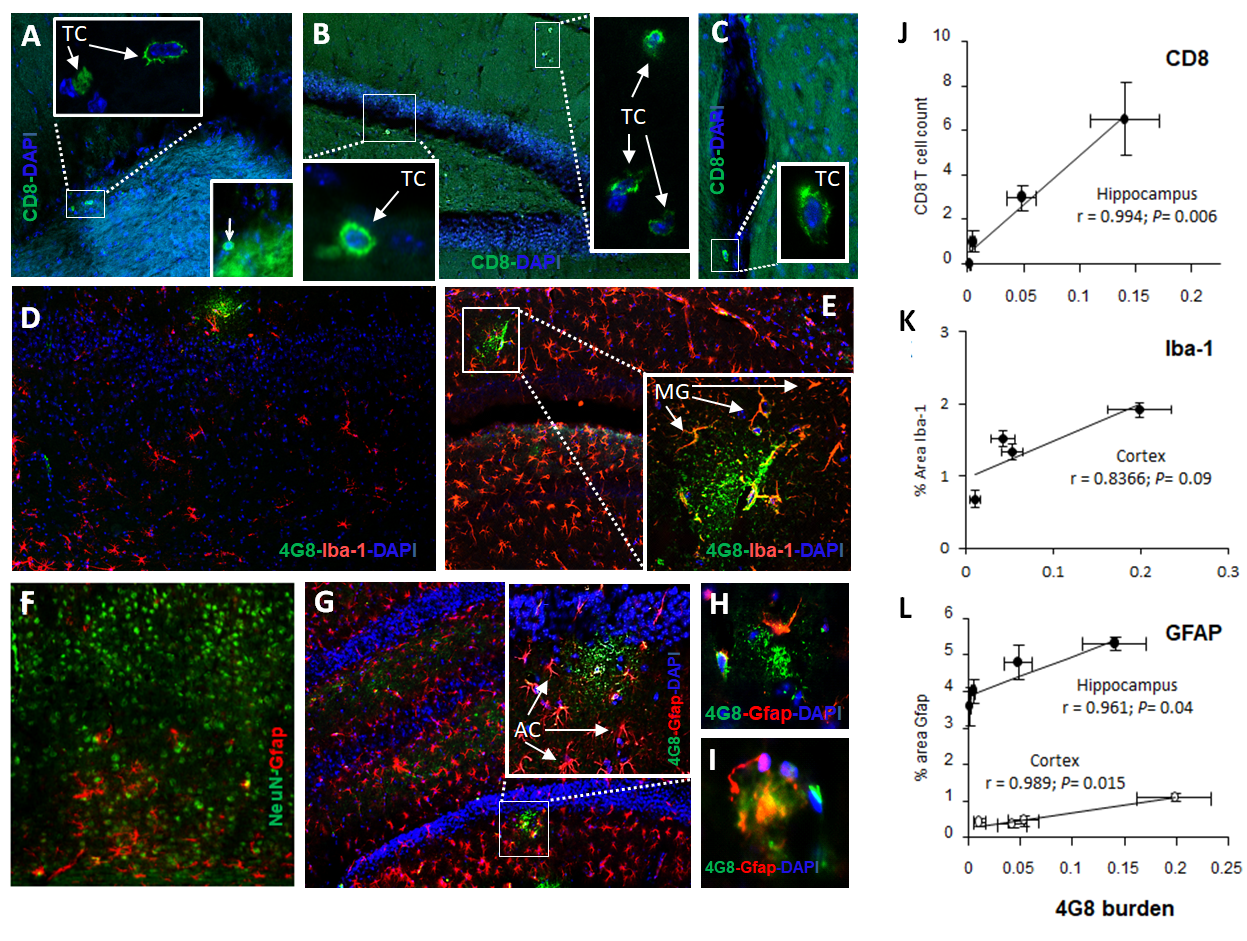

**Fig. S5: Innate and adaptive immune correlates of amyloidosis in nude mice harboring ^hi^T cells.** Brain was co-stained for CD8 and DAPI (**A-C**); Iba-1, 4G8 and DAPI (**D-E**); or GFAP with either NeuN, or 4G8 and DAPI (**F-I**), and quantified within hippocampal and/or cortical brain sections from B6.Foxn1 recipients 15 months after injection of wild-type, IfnγKO or PrfKO CD8 T cells, or PBS as previously reported (Panwar et al., 2020; reference 7 in manuscript). Group data are compiled for CD8 T cell (CD8), microglial (Iba-1), and astrocytic (GFAP) areas and correlated with 4G8^+^ plaque burden within each group, with *P* values of linear regressions and Pearson’s correlations (r) shown (**J-L**). Numbers of mice per group are in Supplemental Table 1.

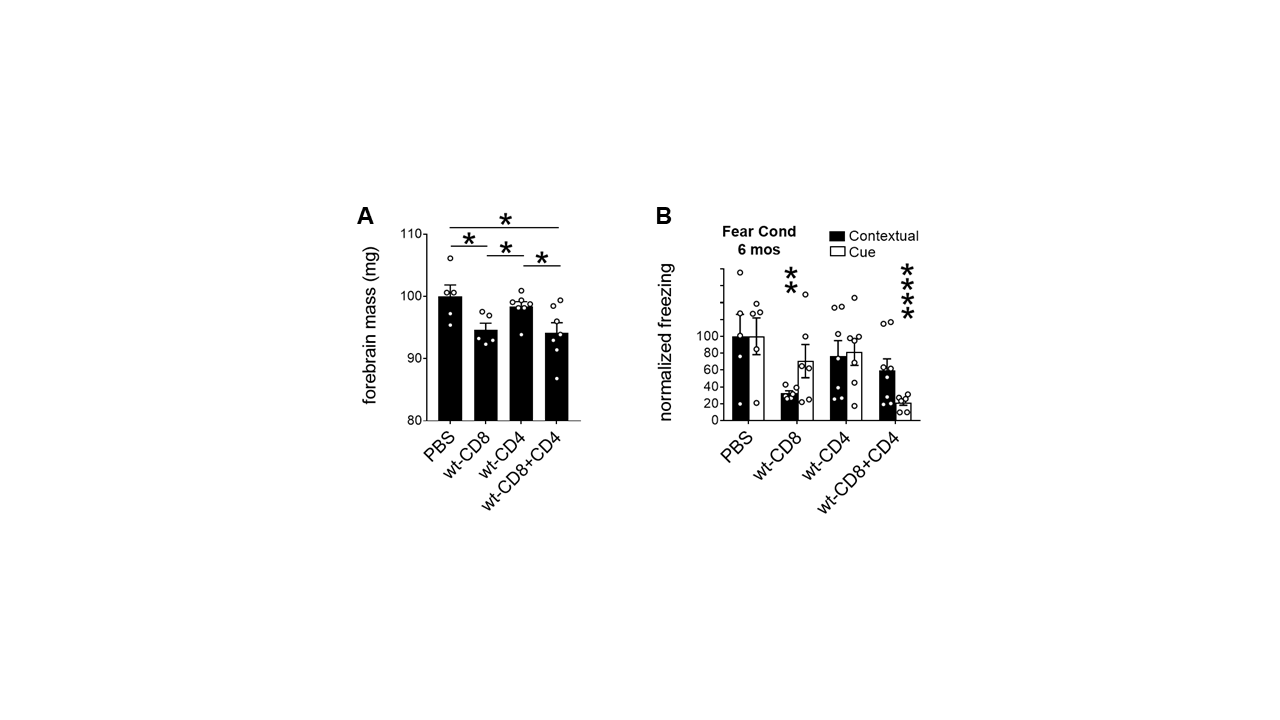

**Fig. S6: Brain mass and behaviour in wt-CD4 and wt-CD8+CD4 recipients.** Brain mass normalized to PBS controls in PBS and wt-CD8 groups (also shown in Fig. 2d), as well as wt-CD4 and wt-CD8+CD4 groups, 6 months after injection (**A**). Fear Conditioning in PBS and wt-CD8 groups (also shown in Fig. 3B), as well as wt-CD4 and wt-CD8+CD4 groups, 6 months post-injection **(B)**. **P* < 0.05, ***P* < 0.01, ****P* < 0.005 by 2-sided T-test relative to PBS group.

**A**

**B**

**C**

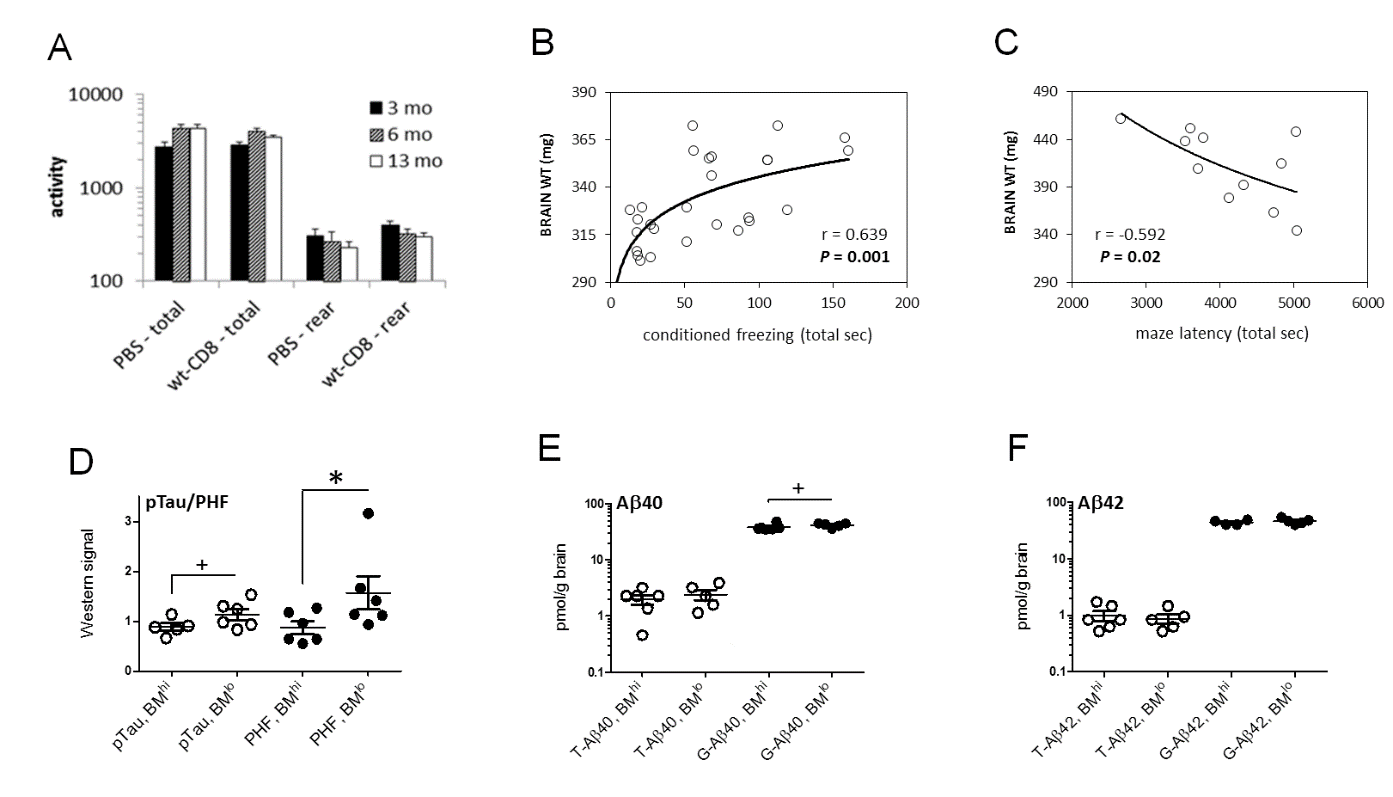

**E**

**F**

**D**

**Fig S7: Motor activity, cognitive performance and correlation with pathological features.** Open Field total activity and rearing activity at 3, 6, and 13 months post-injection of CD8 T cells in experimental mouse groups (**A**). There was no substantial alteration in total or rearing activity between PBS and wt-CD8 groups at any time point, although total activity significantly increased after 3 months, and rearing activity significantly decreased by 13 months, in both groups (n ≥ 9 mice/group). Individual mouse performance in Fear Conditioning test at 6 months correlated directly with brain mass (n = 27; mice were from PBS and wt-CD8 groups; **B**). Superior performance of individual mice in Barnes Maze at 14 months was significantly associated with higher brain mass (n = 9; mice were from all groups; **P* > 0.05, ^+^*P* > 0.1 by 2-tailed T-test; **C**). Stratification by median latency in Barnes Maze correlated significantly with Tau PHF only (**D**), although marginal non-significant trends were observed for pTau and GuanidineHCl-soluble Aβ40 relative to Triton X-100-soluble Aβ species (G-Aβ and T-Aβ, respectively; BM^hi^ = longer latency; BM^lo^ = shorter latency; **E,F**).

**Fig. S8: Doublecortin (DCX)^+^ neurons in ^hi^T mice.** Staining of CD8 and Doublecortin (DCX) in 6 month-old C57Bl/6 brain (**A**), and in PBS, IfnγKO-CD8, and wt-CD8 group brain 15 months after i.v. injection (**B, C**). Main panels of **A** and **B** are at 20x magnification, with inset shown at 10x magnification. Left ventricle (L V) is denoted for positional reference. N ≥ 8 mice/group; ****P* < 0.005 by 2-tailed T-test.

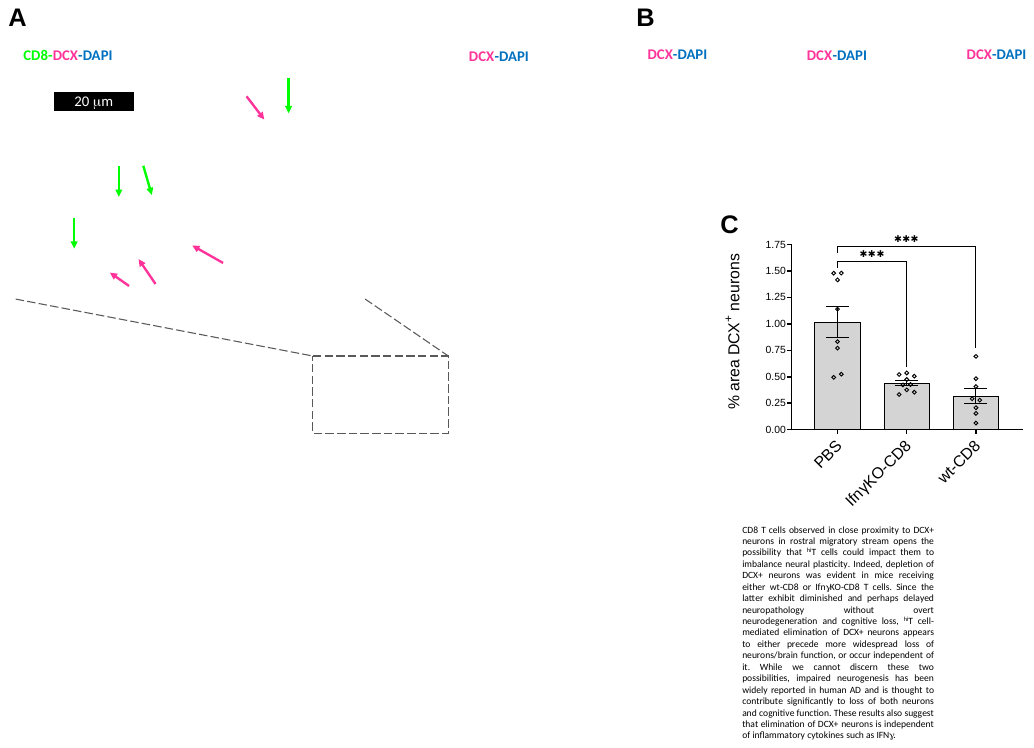

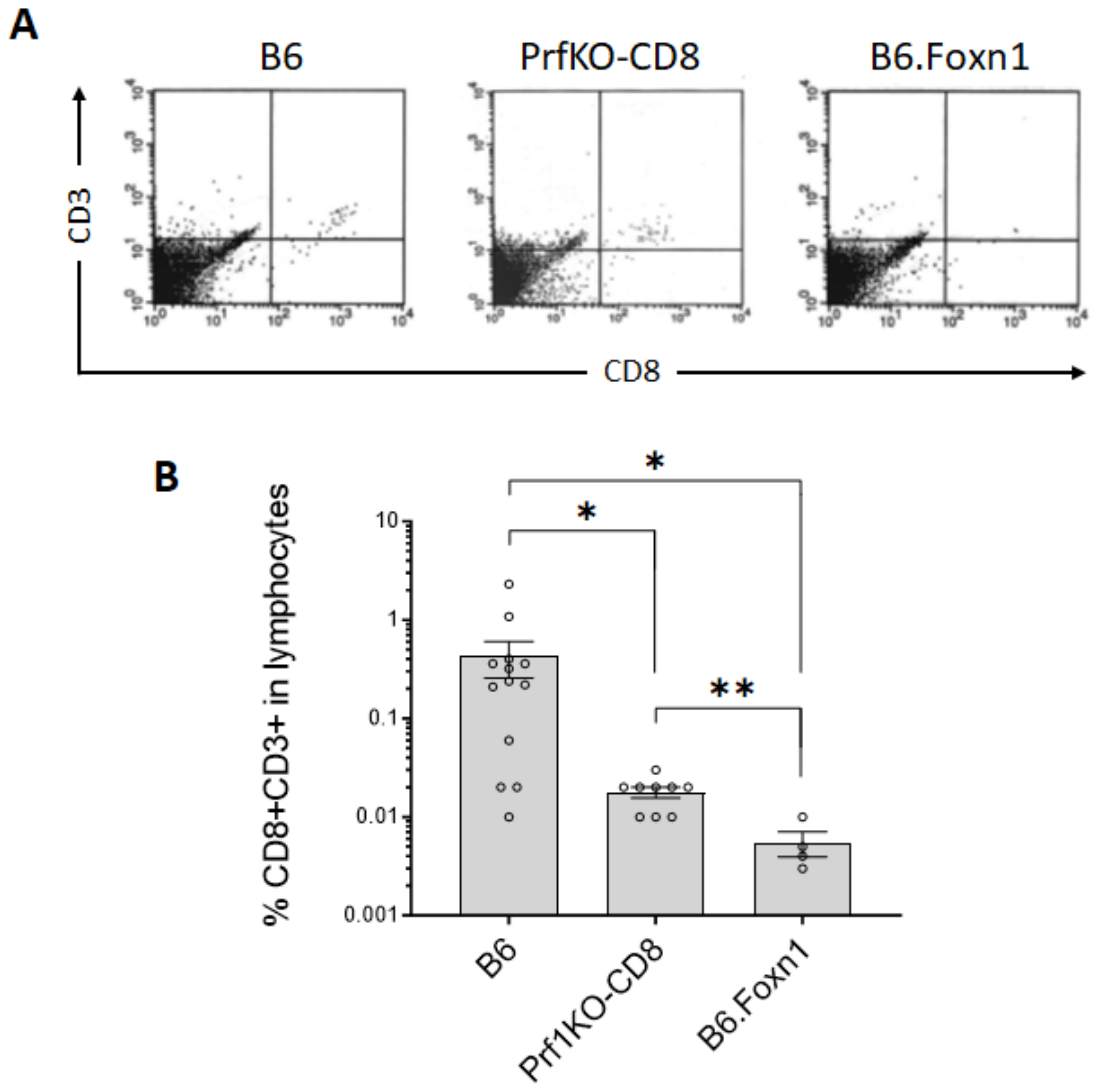

**Fig. S9: Flow cytometric assessment of CD8 T cells in Prf1KO-CD8 group brain.** Staining of CD8 and CD3ε in wild-type C57BL/6 (B6), PrfKO-CD8, and untreated B6.Foxn1 individual forebrains (**A**), with compilations from multiple biological replicates in (**B**). ***P* < 0.01; **P* < 0.05 by 2-tailed T-test.

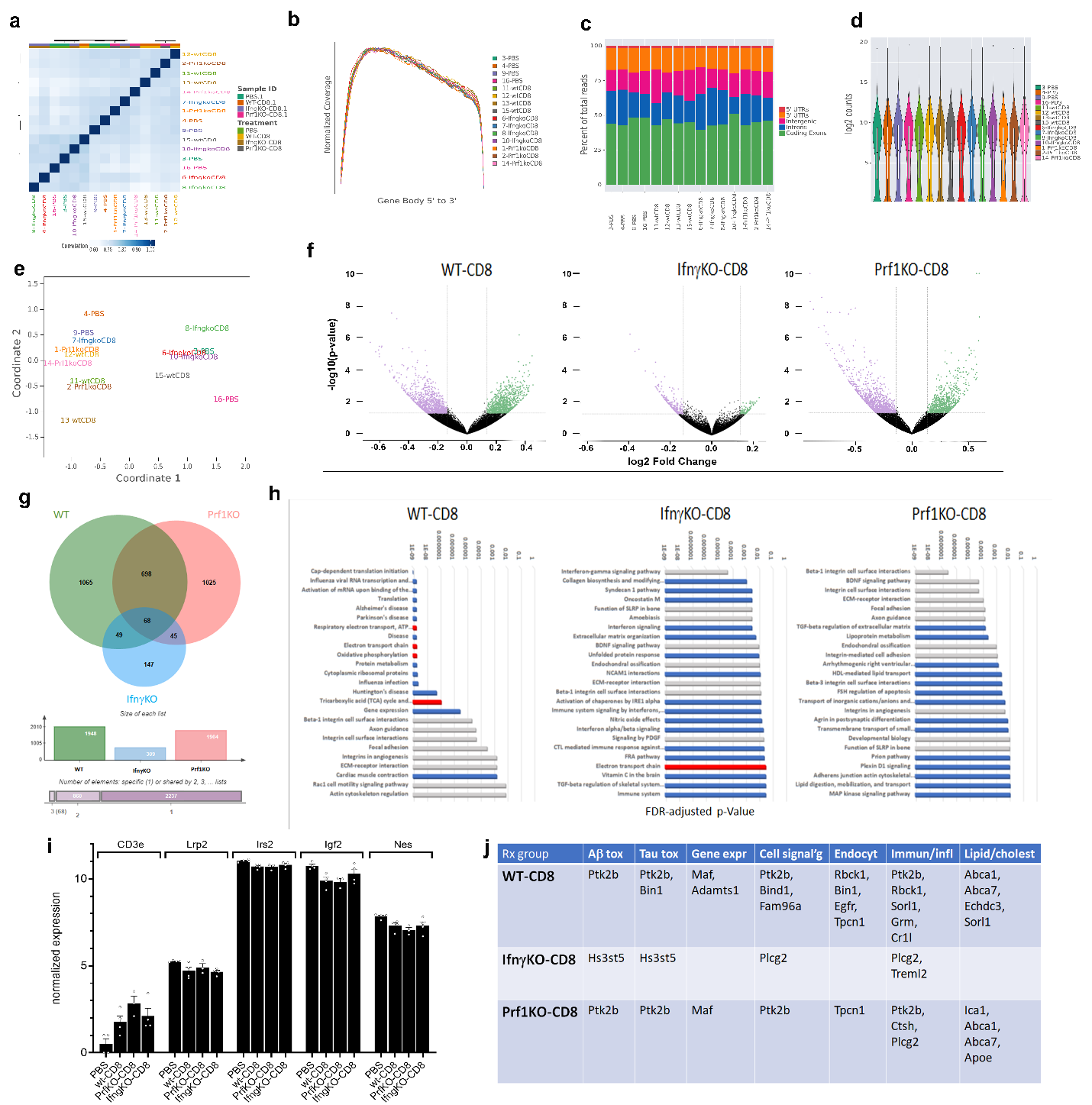

**Fig. S10: Quality control and detailed pathway analysis on RNAseq from brains of ^hi^T mice.** Sample correlation heatmap with data matrix containing correlation values between samples; darkest blue represents the strongest correlations. Dendrogram annotation on top axis indicates clustering of samples (**A**). **Gene Body Bias showing how** mapped reads were distributed over gene bodies, averaged over all expressed genes: 5' end of all genes is represented on the left of graph, with 3' end represented on the right (**B**). **Genomic region distribution** plot shows how mapped reads were distributed over gene features across the genome (**C**). **B**ox plots (base 5 metrics) overlayed on violin plot to show summary statistics, data distribution of the log of the un-normalized gene counts for each sample, and data variation (**D**). Multidimensional scaling (MDS) plot of expression differences between samples within the RNAseq experiment (**E**). Volcano plots of each T cell-treated group relative to PBS controls; X axis = Log ratio of the fold change; Y axis = negative log of p-value (**F**). Each dot represents a gene within the comparison. Venn diagram of numbers of genes differentially regulated in each treatment group, and their overlap, relative to PBS controls (**G**). FDR-adjusted P values of Top 25 BioPlanet pathways significantly altered relative to PBS controls in each T cell-treatment group (**H**). Grayed bars depict pathways present in PrfKO-CD8 and other groups; Red bars depict pathways shared between wt-CD8 and IfngKO-CD8 groups. Normalized levels of CD3e and selected non-GWAS genes relevant to AD in all groups and samples; all shown exhibit P < 0.05 in at least on T cell-treated group, with CD3e *P* < 0.02 in all treatment groups, relative to PBS controls (**I**). Significantly altered AD-associated (“GWAS”) genes in each T cell-treatment group categorized by biological process (**J**).

Fig. S11: KEGG pathway analysis on differentially expressed genes in brains of wt-CD8 group ^hi^T mice. 1943 differentially expressed genes from wt-CD8 group forebrains (n = 4; +/- 1.1 relative to PBS group) were mapped onto the KEGG Alzheimer’s Disease pathways using the DAVID website. Red asterisks denote at least one significantly differentially regulated gene within the affected gene group (*P* < 0.05).

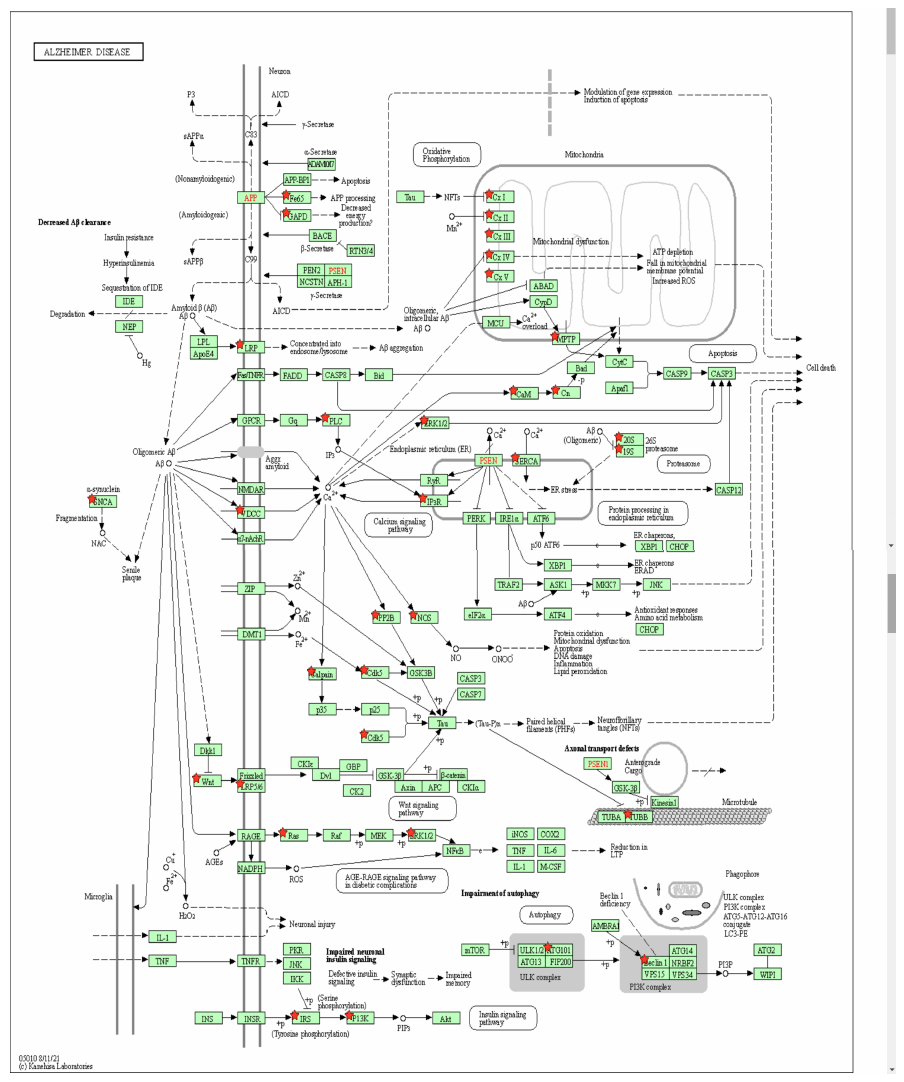

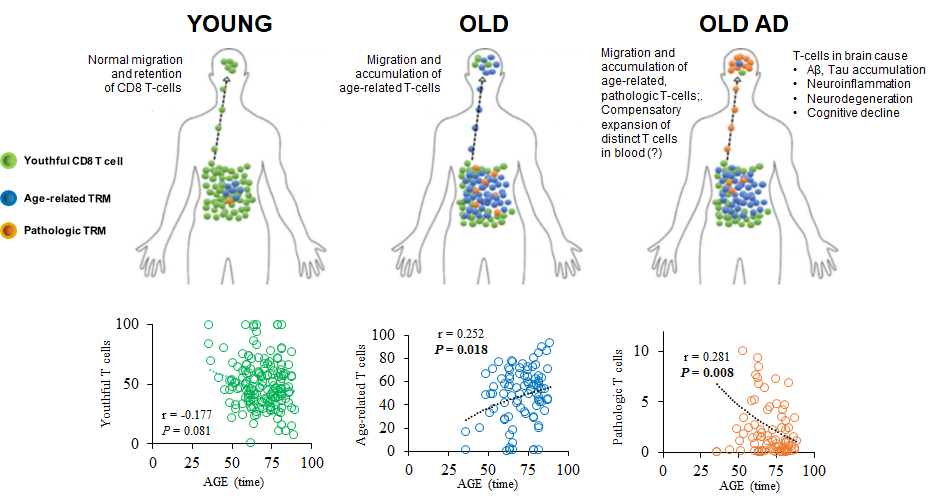

**Fig. S12: KLRG1^-^ (“youthful”), KLRG1^+^ (“age-related”), and APP_(471-479)_/HLA-A2 multimer-reactive KLRG1^+^ (“pathologic”) CD8 T cells exhibit differential associations with age.** Whole blood from 165 human patients (31 MCI – norm bio; 45 MCI – AD bio; 40 normal aging controls; 49 AD) was analysed by flow cytometry for KLRG1^+^ and KLRG1^-^ CD8 T cell content in lymphocyte gates, and plotted relative to patient age at blood collection, with APP_(471-479)_/HLA-A2 multimer reactivity within KLRG1^+^ CD8 T cells further quantified in the subset of 88 HLA-A2^+^ patients. Possible migration of age-related and APP-reactive T cell subsets from blood to brain as depicted above the plots is proposed.

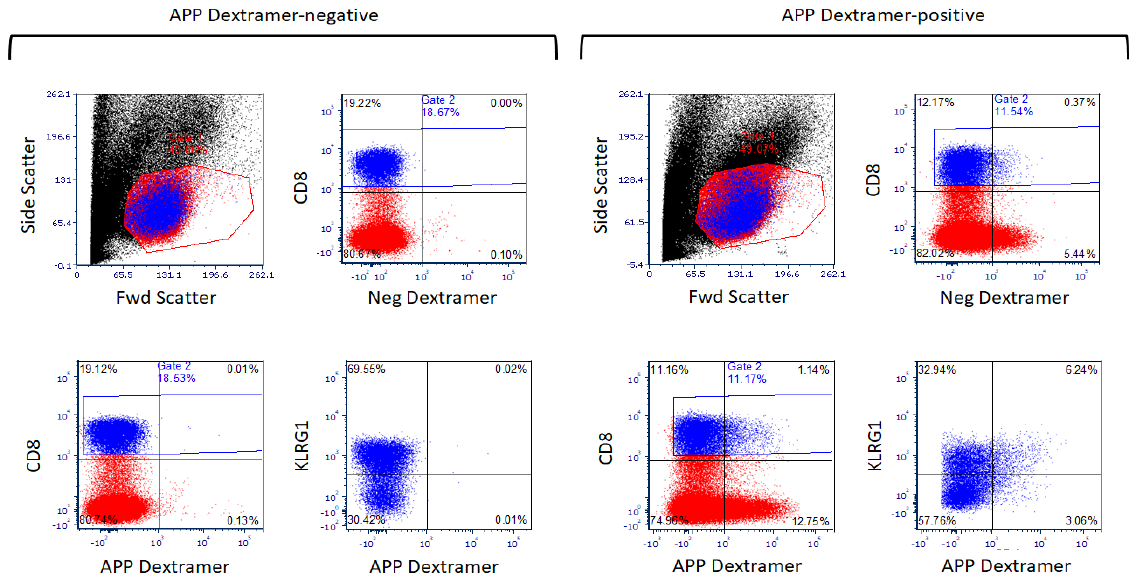

**A**

**B**

**C**

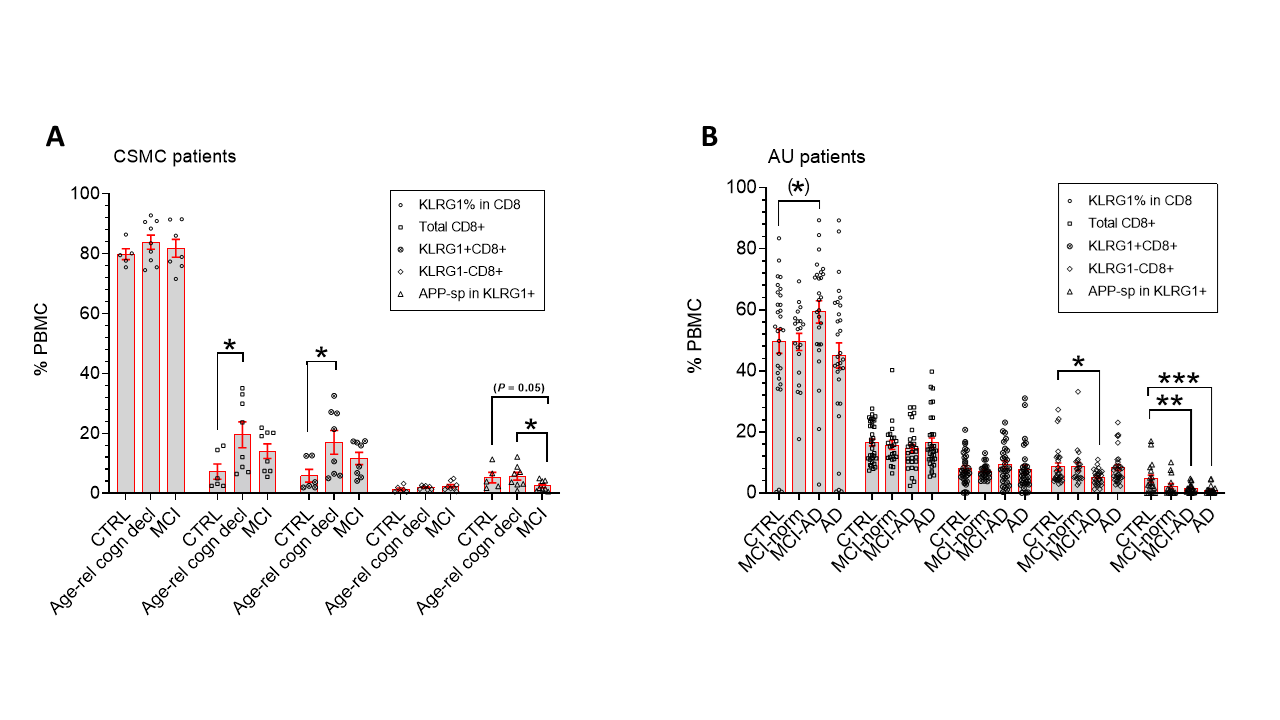

**Fig. S13: Parental KLRG1^+^ and APP-specific KLRG1^+^ CD8 T cell levels two independent cohorts.** Representative flow cytometric plots and gating of APP dextramer positive and negative samples (**A**). KLRG1^+^, KLRG1^-^ and APP_(471-479)_/HLA-A2-reactive KLRG1^+^ CD8 T cell subpopulations in control and MCI patients from Cedars-Sinai Medical Center (CSMC; **B**) and from control, MCI, and AD patients from Antwerp University (AU; **C**). **P* < 0.05, ***P* < 0.01, ****P* < 0.005, *****P* < 0.001 by 2-sided T-test with 1-sided T-test values depicted in parentheses, relative to CTRL unless otherwise indicated.

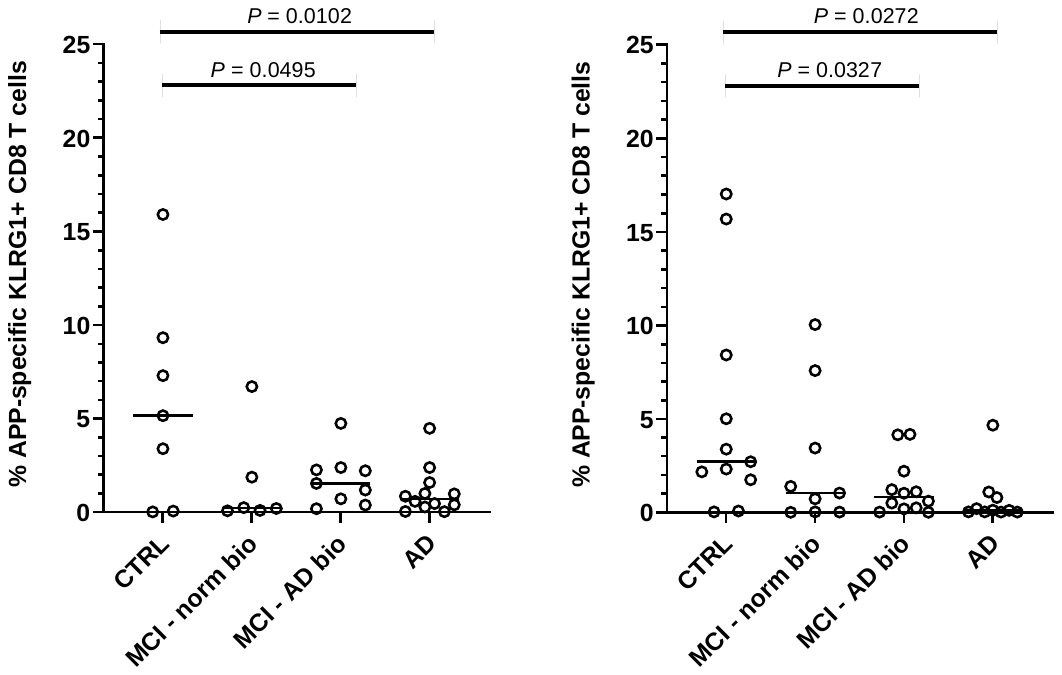

**B**

**A**

**Fig. S14: APP-specific KLRG1^+^ CD8 T cell levels by sex in University of Antwerp cohort.** APP_(471-479)_/HLA-A2-reactive KLRG1^+^ CD8 T cells in males (**A**) and females (**B**) in CTRL, MCI ± CSF AD biomarkers (MCI, MCI–AD), and verified Alzheimer’s (AD) blood. Significance *P*, by 2-sided T-test relative to CTRL.
**Fig. S15: Additional human cohort analysis: increased ^hi^T cell-associated surface markers in human AD brain and blood.** GFAP expression in all-stage (insipient, moderate, severe) AD brain from microarray database (n ≥ 9; GEO accession #s GSM21203-GSM21233)(**A**). Additional indicated genes were normalized to average of GFAP in all AD combined, zeroed on the average of controls for each individual gene, and percent up- or down-regulation relative to GFAP in AD patients (**B**). GFAP expression itself was not significantly increased in insipient AD, but was significantly elevated in moderate, severe, and all AD combined, relative to controls (*P* = 0.1185, 0.0002, 0.0112, and 0.0008, respectively; 1-sided T-test). Relative expression of cytolytic (CD8) resident memory T cell marker genes (CD8A, CD44, CD103; GEO accession # GSE85426) demonstrating significantly decreased CD4, but significantly increased CD8A, CD44, and CD103 expression, consistent with elevated CD8 T_RM_ in AD blood (n≥39 /group; **C**). Other T_RM_ markers were either not available, or were not significantly altered in this dataset. Samples were adjusted for T cell level and age (above or below median expression of CD3D, a pan-T cell marker, and age > 65; **D**), prior to biomarker analysis. **P* < 0.05, ***P* < 0.01, ****P* < 0.005, *****P* < 0.00005, 2-sided T-test.

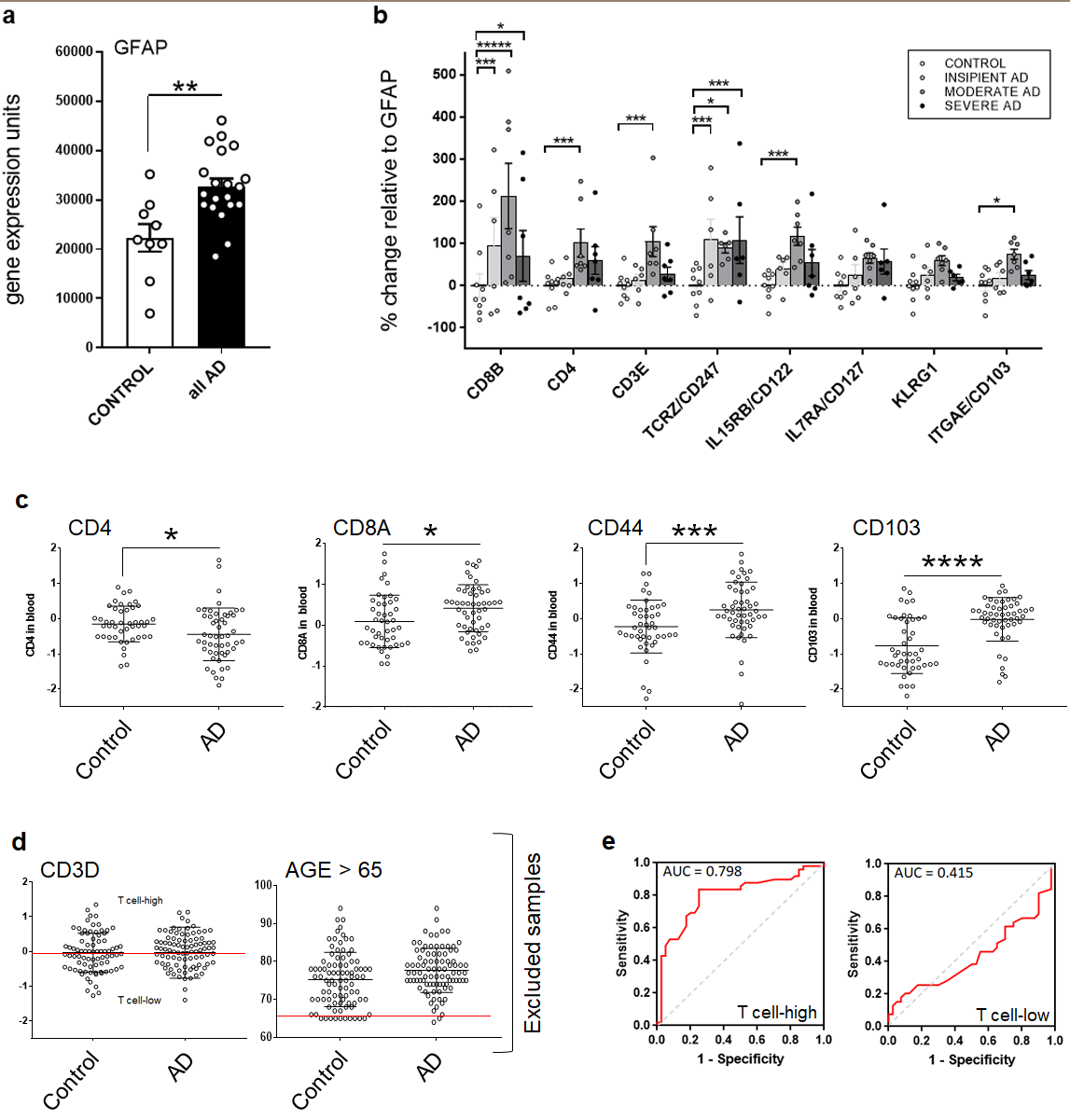

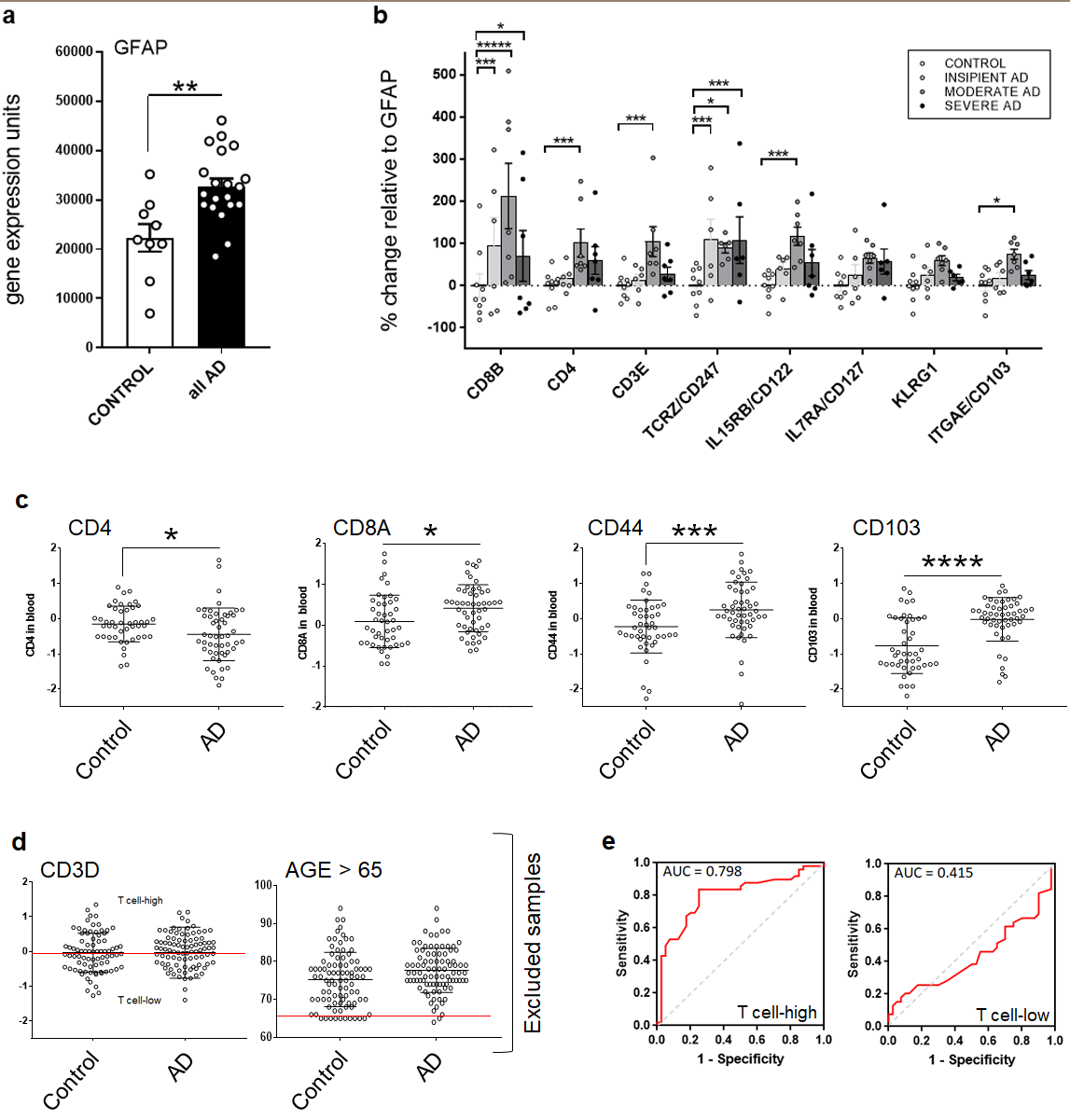

**A**

**B**

**C**

**D**

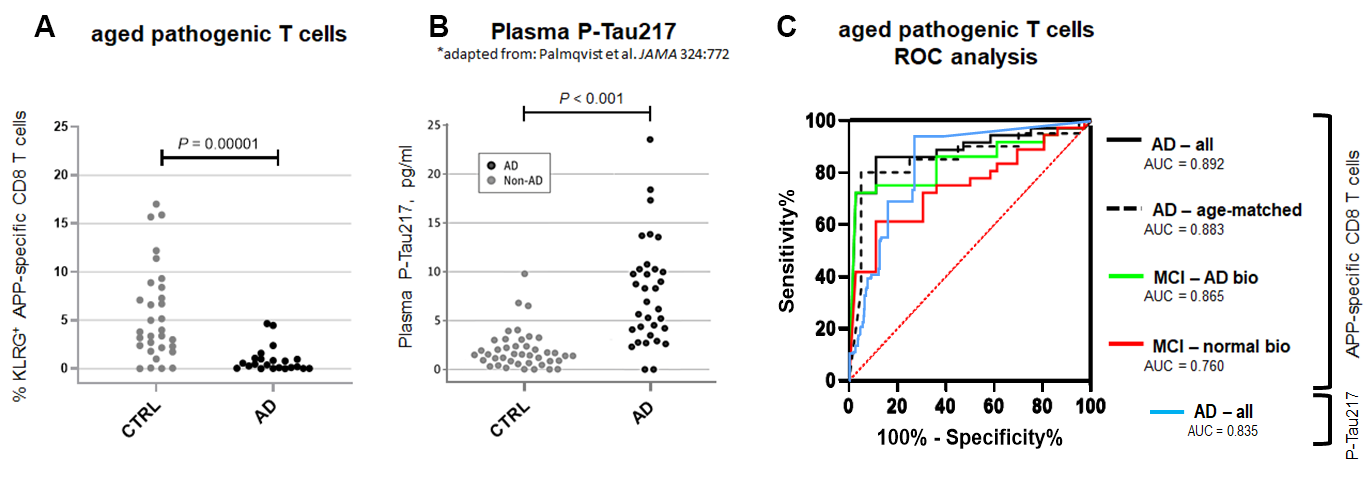

**Fig. S16: Comparison with P-Tau217 and receiving operator characteristic (ROC) curves for APP-specific KLRG1^+^ CD8 T cell level in human patients.** Levels of APP/HLA-A2 multimer-reactive KLRG1^+^ CD8 T cells in blood of AD patients from our study (**A**) relative to published plasma levels of P-Tau217 (Palmqvist et al., 2020. JAMA 324(8):772-781; **B**). Receiver Operating Characteristic (ROC) plots of APP/HLA-A2 multimer-reactive KLRG1^+^ CD8 T cells in blood of indicated patient cohorts relative to normal aging controls (**C**) Mild Cognitive Impairment without (MCI-normal bio) and with (MCI-AD bio) CSF biomarkers consistent with AD, and confirmed AD patients ages 57-84 (AD-all). Area Under the Curve (AUC) is indicated. AD-age-matched indicates ROC analysis on 10 AD patients for whom precisely age-matched controls were available (+/- 1 year; n = 10). *P* < 0.001 for all curves except MCI – normal bio (*P* = 0.003).

Table S1

| **HOST** | **ANALYSIS** | **Brain Region/Method** | **EXPERIMENTAL GROUP “n”** | | | | **VALIDATION** |
| --- | --- | --- | --- | --- | --- | --- | --- |
|  |  |  | **PBS** | **wt-CD8** | **PrfKO-CD8** | **IfnγKO-CD8** |  |
| **B6.Foxn1** | CD8 | Cortex | 10 | 11 | 9 | 6 | WB, IHC morphology (Ref 7) |
|  |  | Hippo | 11 | 12 | 3 | 6 |  |
|  | GFAP | Cortex | 10 | 19 | 10 | 6 | WB, IHC morphology (Ref 7) |
|  |  | Hippo | 9 | 14 | 7 | 6 |  |
|  | Iba1 | Cortex | 3 | 5 | 3 | 5 | IHC morphology  (Ref 7) |
|  |  | Hippo | 4 | 4 | 3 | 3 |  |
|  | Aβ (10 wk) | 1-40 | 4 | 9 | NA | NA | NA |
|  |  | 1-42 | 4 | 9 | NA | NA |  |
|  | Aβ (15 mos) | 1-40 | 4 | 7 | 3 | 3 |  |
|  |  | 1-42 | 8 | 14 | 6 | 6 |  |
|  | 4G8 IHC | Cng ctx | 7 | 11 | 6 | 8 | WB (absorbed), IHC morphology, huAD IHC |
|  |  | Hippo | 6 | 11 | 5 | 8 |  |
|  |  | Ent ctx | 4 | 10 | 6 | 8 |  |
|  | pTau/PHF | 10 wk | 7 | 11 | NA | NA | WB, IHC morphology, huAD IHC |
|  |  | 15 mos | 4 | 7 | 6 | 5 |  |
|  | Gallyas | Cng ctx | 7 | 15 | 5 | 9 | IHC morphology, huAD IHC |
|  |  | Hippo | 7 | 13 | 6 | 19 |  |
|  |  | Ent ctx | 19 | 19 | 5 | 10 |  |
|  | NeuN | WB | 8 | 8 | 6 | 6 | WB, IHC morphology |
|  |  | counts | 4 | 6 | 3 | 3 |  |
|  | Drebrin WB | forebrain | 4 | 4 | 3 | 3 | WB, IHC morphology |
|  | Brain wt | 6 mos | 5 | 5 | NA | NA | NA |
|  |  | 15 mos | 4 | 8 | 7 | 7 |  |
|  | Open Field | 3 mos | 10 | 21 | 10 | 10 |  |
|  |  | 6 mos | 10 | 15 | 10 | 10 |  |
|  |  | 13 mos | 7 | 10 | 10 | 10 |  |
|  | Fear Cond | 6 mos | 8 | 11 | NA | NA |  |
|  |  | 11 mos | 16 | 15 | NA | NA |  |
|  | Spont Alt | 12 mos | 7 | 17 | 10 | 10 |  |
|  | Barnes Maze |  | 14 | 12 | 10 | 9 |  |
|  | | | **normal** | **mild AD** | **severe AD** |  | |
| **Human** | microarray |  | 9 | 13 | 7 |  | GFAP normalization |
|  | Prf1 protein |  | 5 | 4 | 10 |  | WB, huAD IHC |
|  | pHLA/APP  +anti-CD8 |  | 10 | NA | 10 |  | IHC morph, co-staining |

**Table S1: Group numbers and validation**. All “n” refer exclusively to biological replicates. Validation: WB = Western blot; morph = expected morphology obtained on tissue staining; WB(absorb) = expected positive signal by Western blot with negative antigen-absorbed control; huAD = additional expected morphology obtained on brain tissue from human AD patients; co-staining = stained with 2^nd^ cell-type-specific reagent, (anti-CD8 antibody).
